# Diversification and recurrent adaptation of the synaptonemal complex in *Drosophila*

**DOI:** 10.1101/2023.10.20.563324

**Authors:** Rana Zakerzade, Ching-Ho Chang, Kamalakar Chatla, Ananya Krishnapura, Samuel P Appiah, Jacki Zhang, Robert L Unckless, Justin P Blumenstiel, Doris Bachtrog, Kevin H-C Wei

## Abstract

The synaptonemal complex (SC) is a protein-rich structure essential for meiotic recombination and faithful chromosome segregation. Acting like a zipper to paired chromosomes during early prophase, the complex consists of central elements bilaterally tethered by the transverse filaments to the lateral elements anchored on either side to the homologous chromosome axes. Despite being found in most major eukaryotic taxa implying a deeply conserved evolutionary origin, several components of the complex exhibit unusually high rates of sequence turnover. This is puzzlingly exemplified by the SC of Drosophila, where the central elements and transverse filaments display no identifiable homologs outside of the genus. Here, we exhaustively examine the evolutionary history of the SC in *Drosophila* taking a comparative phylogenomic approach with high species density to circumvent obscured homology due to rapid sequence evolution. Contrasting starkly against other genes involved in meiotic chromosome pairing, SC significantly shows elevated rates of coding evolution due to a combination of relaxed constraint and recurrent, widespread positive selection. In particular, the central element *cona* and transverse filament *c(3)G* have diversified through tandem and retro-duplications, repeatedly generating paralogs that likely have novel germline functions. In a striking case of molecular convergence, *c(3)G* paralogs that independently arose in distant lineages evolved under positive selection to have convergent truncations to the protein termini and elevated testes expression. Surprisingly, the expression of SC genes in the germline is exceedingly prone to change suggesting recurrent regulatory evolution which, in many species, resulted in high testes expression even though *Drosophila* males are achiasmic. Overall, our study recapitulates the poor conservation of SC components, and further uncovers that the lack of conservation extends to other modalities including copy number, genomic locale, and germline regulation. Considering the elevated testes expression in many Drosophila species and the common ancestor, we suggest that the function of SC genes in the male germline, while still poorly understood, may be a prime target of constant evolutionary pressures driving repeated adaptations and innovations.

**Summary:** The synaptonemal complex (SC) is essential for meiotic recombination and faithful chromosome segregation across eukaryotes, yet components of the SC are often poorly conserved. Here we show that across the *Drosophila* phylogeny several SC genes have evolved under recurrent positive selection resulting in orthologs that are barely recognizable. This is partly driven duplications repeatedly generating paralogs that may have adopted novel germline functions, often in the testes. Unexpectedly, while most SC genes are thought to be dispensable in the male germline where recombination is absent in *Drosophila*, elevated testes expression appears to be the norm across the genus and likely the ancestral state. The evolutionary lability of SC genes in *Drosophila* is likely a repeated source of adaptive innovations in the germline.

## INTRODUCTION

Meiotic recombination, the exchange of non-sister, homologous chromosomes through physical crossovers, is an essential genetic mechanism universal to sexually reproducing eukaryotes. It allows for the shuffling of homologous alleles generating novel allelic combinations. This is necessary for maintaining nucleotide diversity and efficacy of selection; without it, chromosomes (like on the non-recombining, degenerate Y or W chromosomes) will irreversibly accumulate deleterious mutations ultimately leading populations to go extinct. At the cellular level, meiotic pairing, synapsis, and resolution of double strand breaks into crossovers are critical for stabilizing meiotic bivalents as failure is typically associated with skyrocketing aneuploidy rates. Therefore, recombination is a crucial genetic process that is necessary for reproductive fitness and long-term species survival.

Despite the critical functionality of recombination and the deep conservation across eukaryotes, aspects of this fundamental genetic mechanism are surprisingly prone to change. Recombination rate has been repeatedly shown to vary drastically between closely related species [1]. Adaptive explanations typically invoke changing environmental (e.g. temperature [2]) or genomic conditions (e.g. repeat content [3]) requiring commensurate shifts in recombination rate to maintain fitness optima [4,5]. Others have suggested intragenomic conflicts with selfish elements [6] or sexual conflict creating unstable equilibria for optimal fitness [7,8]. However, some have argued that changes in recombination rate have little impact on fitness and rate changes are the byproduct of selection on other aspects of the meiotic processes [9]. Several key findings supporting the adaptive interpretation come from *Drosophila* as multiple genes in the pathways necessary for recombination show signatures of rapid evolution due to positive selection [10–13]. Moreover, because recombination is absent in *Drosophila* males and the SC does not assemble during spermatogenesis [14], sexual antagonism due to sex-specific optima of crossover rates is unlikely the underlying driver of adaptive recombination evolution, at least in species with sex-specific achiasmy. Why recombination, an essential genetic mechanism, is prone to change and whether such changes are adaptive remain central questions in evolutionary genetics [15–17].

The paradox of poor conservation but crucial function is exemplified by the synaptonemal complex (SC), a crucial machinery necessary for meiotic recombination in plants, animals, and major lineages of fungi [18]. It is a protein complex that acts as zippers to tether homologs together along the chromosome axes during meiotic prophase I and forms train track-like structures which have been visualized under electron microscopy across eukaryotic taxa [19,20]. The SC is mirrored along a central axis composed of central element proteins that are tethered by the transverse filaments to lateral elements on two sides anchoring into chromatin (Figure 1A) [21]. This highly stereotypical configuration is found in baker’s yeast, mice, flies, and plants, indicative of an evolutionary ancient structure. Yet, despite the deep evolutionary origin and functional necessity across wide eukaryotic domains, there are many examples of unexpected exceptions. At the extreme are recombining species such as the fission yeast that entirely forego the SC [22]. In another instance, the SC of *Caenorhabditis* has been reconfigured such that the transverse filament – typically a single gene in most SCs – is composed of at least four genes [23]. Therefore, parts of the SC appear to be curiously flexible in composition whereby different analogous but perhaps non-homologous pieces can be recruited and replaced [24].

**Figure 1.**
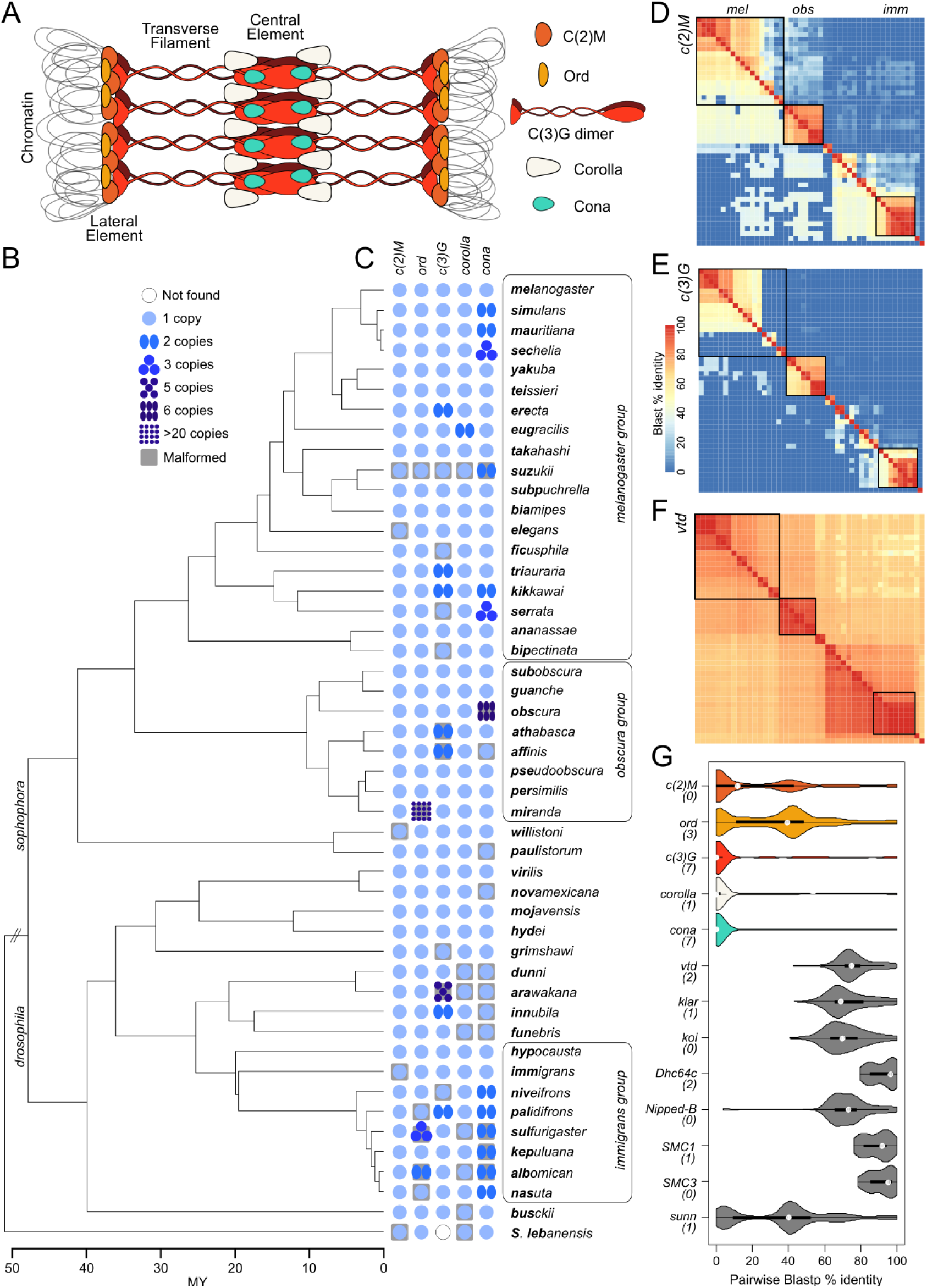
Sequence conservation, or the lack thereof, of synaptonemal complex components across the *Drosophila* genus. A. Cartoon diagram of the Drosophila SC and its primary constituents. B. Phylogenetic relationships of the 48 species used in this study. Bold characters in the species name denote species shorthands. Species dense groups are labeled and boxed. C. Presence, copy number, and absence of SC components across the phylogeny. The number of blue circles indicated the copy number. D-F. Pairwise blast sequence alignments between orthologs from representative species across the genus. Alignments above the diagonal are from nucleotide blasts of the CDS sequences using blastn. Alignments below the diagonal are from protein blasts of the amino acid sequence using blastp. % blast identity is the length of blast alignments multiplied by the % sequence identity, summed across the gene. G. Distribution of pairwise blastp % identity across the genus for SC (colored) and genes (gray) involved in meiotic pairing during early prophase.

Consistent with this flexibility, sequences of SC components are often poorly conserved at shorter evolutionary time scales [12,25]. In *Drosophila*, positive selection appears to repeatedly drive the sequence evolution of the SC, which is composed of the central elements *corona* (*cona*) [26] and *corolla* [27], the transverse filament *c(3)G* [28], and the lateral elements *orientation disruptor* (*ord*) [29] and *c(2)M* [30]. Previously, orthologs of the central region components, *corolla*, *cona,* and *c(3)G,* could not be found outside of the *Drosophila* genus [12] either reflecting divergence so extensive that orthology is no longer recognizable, or novel acquisitions of SC components. Flexibility in SC composition may explain how these molecular transitions are possible without major fitness impacts, but cannot account for why SC genes appear to be evolving under recurrent adaptation. The recent explosion of high quality *Drosophila* species genome assemblies [31–37] offer a unique opportunity to understand the genetic and evolutionary mechanisms driving the strikingly rapid divergence of SC genes. Here, we systematically revisit the evolution of the SC in *Drosophila* by examining the genomes and transcriptomes of 48 species scattered across the entire *Drosophila* phylogeny, with dense representation from three key species groups (*melanogaster*, *obscura*, and *immigrans*). In our exhaustive analyses, we uncovered frequent duplications of several SC components generating paralogs with likely novel functions, in addition to repeated sequence evolution under positive selection. Further, we revealed unexpectedly high rates of expression divergence and regulatory turnover in not just the ovary but also the male germline, where SC genes are thought to have no function. In fact, testes-biased expression of SC genes appears to be the norm, and likely the ancestral state, suggesting SC components have crucial function in male germline, despite the absence of male recombination. Altogether our study revealed a highly dynamic evolutionary history with repeated bouts of copy number, sequence, and regulatory evolution that contribute to the overall poor conservation of SC genes. Further, the surprising transcriptional activity of SC genes in the male germline raises new possibilities for functions of SC genes unrelated to recombination under repeated directional selection in addition to their roles in chiasmate meiosis in the female germline.

## RESULTS

### Poor sequence conservation and frequent duplications of components of the SC

To identify *Drosophila* SC homologs we elected to focus on only species with high quality genome assemblies with either available annotations and/or RNA-seq data (Supplementary table 1). In addition, we strategically generated highly contiguous assemblies of two additional species (*D. hypocausta* and *D. niveifrons*, belonging to the immigrans group; Supplementary table 2), and testes and ovaries RNA-seq of eight species (*D. subobscura*, *D. arawakana*, *D. dunni*, *D. innubila*, *D. funebris*, *D. immigrans*, *D. hypocausta*, and *D. niveifrons*) to either annotate previously unannotated genomes or to refine previous annotations (Supplementary table 1). Altogether, we compiled a total of 47 species spanning the two major arms of the *Drosophila* genus (the *Sophophora* and *Drosophila* subgenera), with three species groups particularly well-represented (*melanogaster*, *obscura*, and *immigrans* groups) (Figure1B) and the outgroup species *Scaptopdrosophila lebanonensis*.

Using a multi-step reciprocal best blast hit approach, we sought to identify orthologs and paralogs across the genus (Figure 1B and C; Material and methods). However, we found that the gene structures are frequently malformed regardless of the source of the annotation (publicly available, or our ones we generated). SC componenets are often mis-annotated as truncated or chimeric gene products or entirely missing in the annotation (Supplementary table 3; for examples see Supplementary figure 1), likely due to the combination of exacerbating factors such as poor sequence conservation, frequent presence of tandem duplicates, low RNA-seq reads, and in some cases assembly errors. To ensure proper sequence alignments, we therefore curated all SC genes and manually re-annotated all erroneous ones ensuring at a minimum, well-formed CDSs and intact full ORFs (Supplementary figure 1 and 2; see Materials and Methods). Note, because *cona* is a short gene with few exons, we hand-annotated its orthologs in 8 additional species (see below).

For the lateral elements *c(2)M* and *ord*, sequence homology is somewhat preserved across the genus, although homology is not always detectable between distant species using Blast (Figure 1D and Supplementary Figure 3A). However, for the central region genes (*c3G*, *cona*, *corolla*), DNA sequence homology quickly becomes unrecognizable outside of species groups, while weak protein homology is only occasionally detectable (Figure 1E, Supplementary Figure 3A). Previously, Hemmer and Blumentiel 2018 identified SC orthologs in a subset of fly species [12]. Increased species and better annotations enabled us to identify orthologs previously missed (*cona* in *D. willistoni* and *corolla* in the outgroup) and resolved discrepant homology relationships (*cona* in the *Drosophila* subgenus, see below). To determine whether this lack of conservation is common to other genes involved in early meiotic progression, we curated and identified the homologs of 8 genes necessary for meiotic chromosome pairing (Figure 1F and Supplementary Figure 3B) across the *Drosophila* genus. The poor sequence conservation of the SC starkly contrasts from these (Figure 1G): even the most conserved component of the SC, *ord,* shows significantly poorer sequence homology compared to the least conserved meiotic pairing gene, *sunn* (Figure 1G; Wilcoxon’s Rank sum test p = 1.813e-05).

Furthermore, we revealed multiple independent duplication events, with *c(2)M* being the only SC gene that remained single-copy across the genus. All SC paralogs were previously unaccounted with the only exception being *ord* duplicates in *D. miranda*, which was identified to have rampantly amplified creating over 20 copies (Supplementary figure 2) on the specie’s unique neo-sex chromosomes [38]. Of the poorly conserved components, *c(3)G* and *cona* in particular have recurrent copy number changes, having more than two copies in eight and thirteen species, respectively, as the result of independent duplications in at least seven lineages each. This propensity to duplicate is epitomized by the five *c(3)G* in *D. arawakana* and six *cona* copies in *D. obscura*.

### History of repeated duplication and loss of paralogs

Based on protein trees of the SC components, we find that the current copy number distributions reflect at least 3, 7, 1, and 7 independent duplications of *ord*, *c(3)G*, *corolla*, and *cona*, respectively. The majority of paralogs are recent species-specific duplications resulting in short branch lengths (Figure 2A and 3A, and supplementary figures 3). The genomic locations of the copies further reveal that tandem duplications account for the majority of the observed copies. For the transverse filament *c(3)G*, four of the seven duplication events are tandems (Figure 2A,C-D), three of which (including the five copies in *D. arawakana)* are recent and species-specific. In the lineage leading up to *D. athabasca* and *affinis*, an older tandem duplication generated duplicates of *c(3)G* and the neighboring gene *pon*, (Figure 2D); one of the copies which we designated *c(3)G2* is shorter, while showing poorer conservation and longer branch lengths between the orthologs, suggestive of functional divergence (see below). Similarly, for *cona*, three of the seven duplication events are tandems, including the 6 copies in *D. obscura* (Figure 2B, Supplementary Figure 5).

**Figure 2.**
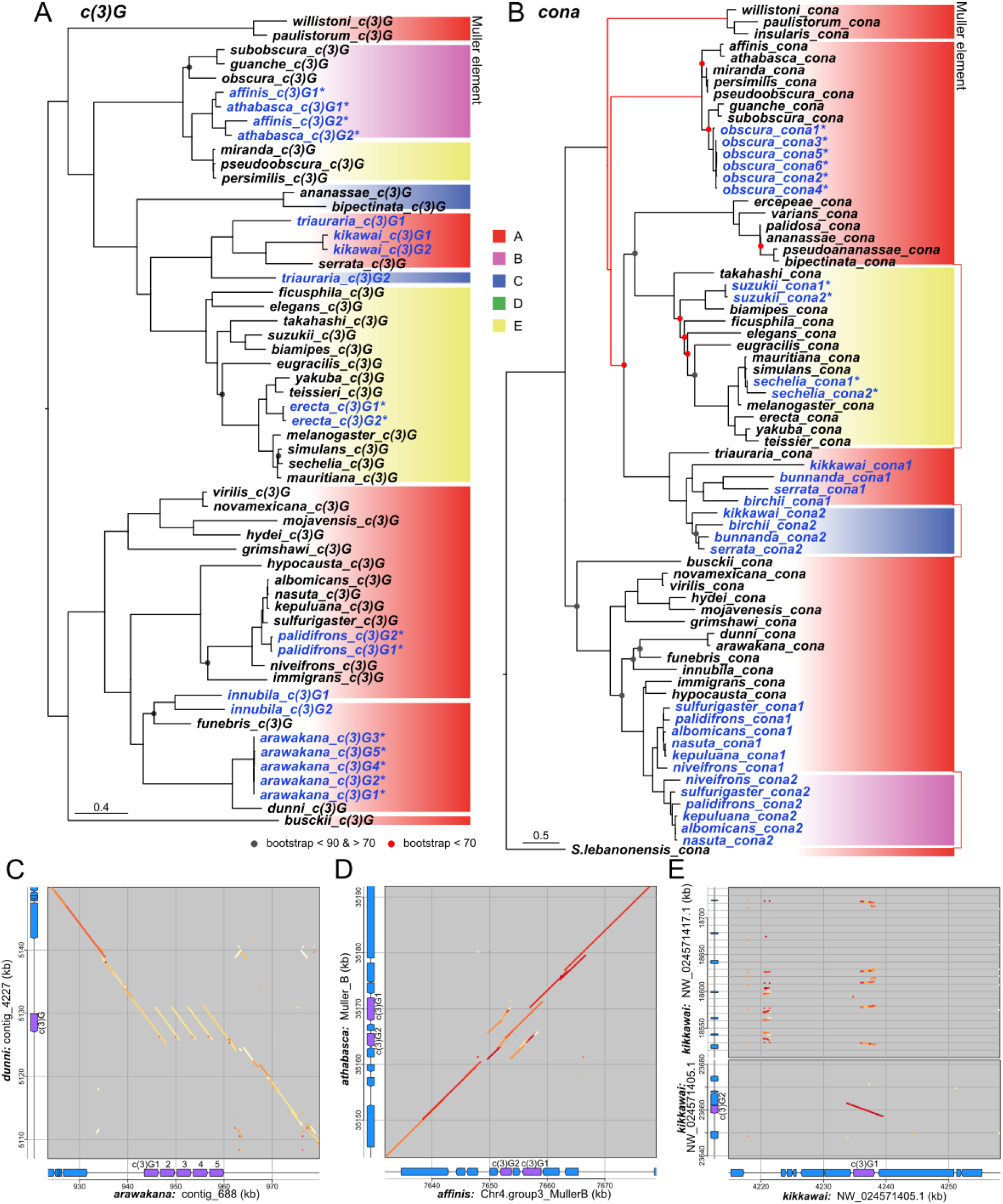
Complex evolution history of synaptonemal complex. A and B. Gene trees of *c(3)G* and *cona* orthologs and paralogs, respectively; nodes with poor bootstrap support are demarcated by circles. Duplicates are labeled in blue and tandem duplicates have asterisks. For *cona*, some branches were adjusted (red) to align with the species tree. For other SC components see Supplementary Figure 4. Color blocks to the right represent different Muller elements on which the genes reside. Unless connected by lines, separate blocks of the same color represent different locations on the same element. C-E. Dotplots showing synteny of genomic regions surrounding *c(3)G* between sister species and/or paralogs. The color of the dots represent the % sequence identity from blastn alignments with darker red reflecting higher identity. In the gene tracks of the displayed regions, *c(3)G* is in purple and other neighboring genes in blue. For E, dotplots depict the local homology of *c(3)G1* and retroduplicate *c(3)G2* (bottom), and additional regions with truncated *c(3)G* (top) in *D. kikkawai*.

**Figure 3.**
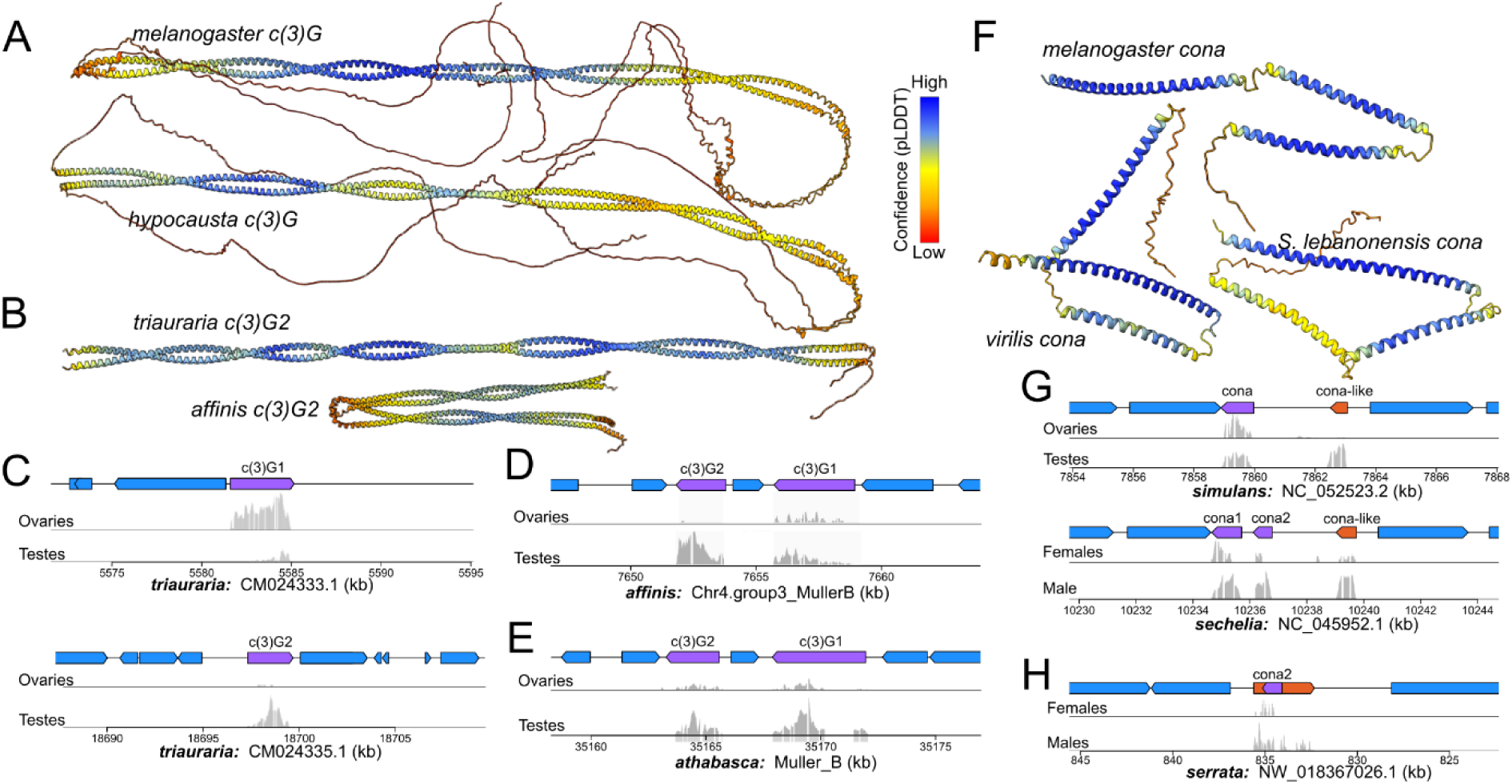
Functional and structural evolution of *c(3)G* and *cona*. A-B. AlphaFold structural prediction of full length c(3)G (A) and diverged paralogs (B). Color represents the confidence of structural prediction. C-E. Gene structure and gonad expression of *c(3)G paralogs* (purple genes) in *D. triauraria* (C), *affinis* (D) and *athabasca* (E). F. Alphafold of distant *cona* orthologs in Drosophila and outgroup species. G. Gene structure and gonad expression of *cona* paralogs in *D. simulans* and *sechelia*. Annotated lncRNA of gene with homology to cona is in orange in the gene tracks. G. Same as F but for D. *serrata*.

Both *c(3)G* and *cona* experienced several instances of likely retroduplications. *cona* offers two clear examples of old events in the common ancestor of the *serrata* and *nasuta* subgroups leading to paralogs shared across all species in the subgroups (Figure 2B). *c(3)G*’s duplication history appears more convoluted but offers unique insight into its dynamic evolution. There are three non-tandem duplicates of *c(3)G* found in *D. kikkawai*, *triauraria*, and *innubila* (Figure 2A); the resulting paralogs found on different chromosomal regions (Figure 2A, Supplementary figure 6A) have minimal neighboring homology to the original (Figure 2E, Supplementary figure 6B) suggesting retroduplication events instead of larger scale duplications. In the latter two species, the duplications are older events evidenced by their phylogenetic placements with long branches separating the paralogs, compared to the species-specific duplication in *D. kikkawai*. For *D. triauraria*, the duplication creating *c(3)G2* predated the split of the serrata species subgroup, but is no longer found in the derived lineages, indicating subsequent loss of the paralog. For *D. innubila,* the paralogs are found 180kb apart on the X chromosome (Supplementary Figure 5B), and the phylogeny suggests that the duplication occurred after the split from *D. funebris*, although with low bootstrap support (Figure 2A). Interestingly, synteny information suggests this is not the true relationship; while one of the *c(3)G* copies is found in the same syntenic block shared with *D. funebris, arawakana*, and *dunni*, the other is in a separate synteny block shared will nearly all other species in the *Drosophila* subgenus and thus the ancestral copy (Figure 2A, Supplemental Figure 6C). This synteny pattern is therefore more parsimonious with an old duplication in the last common ancestor of the dunni, quinaria, and funebris species groups with the original copy lost in species other than *D. innubila*. Non-allelic gene conversion subsequently homogenized the duplicates in *D. innubila*, obscuring the true phylogenetic relationship.

In addition to the one *c(3)G* retroduplicate in *D. kikkawai*, we curiously identified numerous loci across the genome as 5’ truncated homologs, none of which were annotated or have RNA-seq reads mapping (Figure 2E). These truncated and nonfunctional duplicates, along with the two loss events mentioned, raise the possibility that *c(3)G* experienced not only repeated duplications through transpositions, but also repeated pseudogenization events. A similar pattern of nonfunctional duplicates is also observed with *corolla* in *D. arawakana*; despite only one full length *corolla*, there are four adjacent tandem copies that lack the 5’ exon and therefore likely non-functioning (Supplementary figures 5). Further examining the syntenic relationships of the SC homologs, we find that while the lateral elements have maintained the same local microsynteny showing a lack of gene movement, the central region genes have repeatedly relocated to different chromosomes, or different locations on the same chromosome (Figure 2A, Supplementary figure 6A). The X is likely the ancestral home for all three but we documented eleven, sex, and five inter- and intrachromosomal movements for *c(3)G*, *corolla*, and *cona*, respectively. Such recurrent movements likely through transpositions are unusual for flies as chromosomal gene content and microsyteny are largely stable while broad chromosome-scale synteny is scrambled by large scale rearrangements [39]. Curiously, one relatively recent relocation occurred in the common ancestor of the *pseudoobscura* species, moving *c(3)G* from Muller B to an euchromatic repeat block on Muller E (Supplementary figure 6), suggesting that such movements may be mediated by the instability of repetitive sequences, perhaps piggybacking off of transpositions of transposable elements. Since most of these movements no longer have extant paralogs, corresponding pseudogenization events were likely common. Therefore, even though most observable paralogs are young tandem duplicates, retroduplications and pseudogenization events have frequently occurred for *c(3)G, cona,* and even *corolla* which has few remnants of duplicates, thus accounting for the existence of many recent and species-specific copies in different locations but fewer old, shared duplicates.

### *c(3)G* and *cona* paralogs likely have novel germline functions

Much like other transverse filaments, *C(3)G* has an extensive coiled-coil domain flanked by globular domains at the N- and C-termini connecting the central and lateral elements, respectively [18,40]. Despite the poor sequence conservation, we find that this canonical structure is conserved across the genus based on AlphaFold protein prediction (Figure 3A) [41,42] and coiled-coil predictions (Supplementary figure 9) [43]. This unique evolutionary property of structural but not sequence conservation is also observed by Kursel et al. in *Caenorhabditis*, whereby central element genes have conserved coiled-coil domains and near invariant protein lengths, but neutrally evolving sequences [25]. In *D. athabasca*, *affinis*, and *triauraria*, while *c(3)G1*s produce longer proteins (690, 692, 830 AAs respectively) predicted to have the canonical structure, the paralogs *c(3)G2s* all produce notably shorter proteins (361, 395, and 319 AAs, respectively) with the flanking globular domains truncated, if not entirely absent, suggesting they no longer function as transverse filaments that can tether the SC (Figure 3B). Curiously, these paralogs are highly expressed in the testes but lowly expressed in the ovaries (Figure 2C-E), incongruent with the expectation of female meiotic function. Despite being independent duplications in lineages separated by over 25 million years, *c(3)G2* in *D. triauraria* and *D. athabasca/affinis* display remarkably similar structural and regulatory evolution, revealing molecular convergence for likely male germline function. In a single nuclei RNA-seq dataset of *D. affinis* testes, we further find that while *c(3)G1* and *c(3)G2* are both testis-expressed, they are most active in different cell populations (Supplementary Figure 10), evidence that the two have functionally diverged.

Similar to *c(3)G*, *cona* has maintained the same conserved tri-coil structure (Figure 3F), despite poor protein homology. Curiously, several *cona* duplicates similarly show properties that deviate from its characterized function in SC formation during female meiosis. At least two recent duplication events occurred within the *simulans* clade generating two upstream paralogs, one ancestral to the three *simulans* species while another found only in *D. sechelia* (Figure 2B and 3G). The *sechelia*-specific duplicate generates a complete but short ORF and likely protein coding, but the shared paralog only shows homology at the 3’, lacks a complete ORF, and is annotated as a long non-coding RNA (Figure 3G). To evaluate whether this paralog, which we named *cona-like*, is transcriptionally active or pseudogenized, we examined RNA-seq data, and found high expression in the testes and males but low-to-no expression in ovaries or females across all *simulans* species (Figure 3G), strongly suggesting testes function as a lncRNA. Adding to the intrigue, this is not the only instance of a *cona* paralog generating lncRNA. In the *serrata* group, the retroduplicate, *cona2*, is shared across the species (Figure 2B), but only in *D. serrata* does it generate a lncRNA (Figure 2H). Unlike *cona-like* in the simulans clade, this paralog has a well-formed ORF and is expressed in both sexes, suggesting high protein coding potential. However, the lncRNA is anti-sense as confirmed with strand-specific RNA-seq (Supplementary figure 11) and includes additional flanking sequences that only show expression in males. *cona2* likely generates a functional protein in females and ovaries but was incorporated in the anti-sense direction into lncRNA production in the testes of *D. serrata*. Altogether, these results suggest that both *c(3)G* and *cona* paralogs have repeatedly adopted germline functions in the testes unrelated to SC formation.

### *coronetta* (*conta*) is an ancient testes-expressed paralog of *cona*

Previously, Hemmer and Blumenstiel identified *cona* homologs in the *Drosophila* subgenus that were highly diverged from *Sophophora cona* [12]. One of the proteins they identified through reciprocal best blastp hit was *GJ20698* in *D. virilis,* a gene producing a short peptide of 109 AA. We were able to find the orthologs of *GJ20698* across the *Drosophila* subgenus as well as the outgroup, but none of them were reciprocal best hits to *sophophora cona* – in fact, they have no identifiable *sophophara* homologs at all. Instead, increased density of species enabled us to correctly identify *D. virilis’s GJ16397* (the *D. virilis’s* 2nd best hit to *sophophora cona*) which has orthologs across the *Drosophila* subgenus and *S. lebanonensis* that are reciprocals best hits with *sophophora cona* (Figure 1F, 2B). The gene tree affirms that *sophophora cona* is more closely related to *GJ16397,* which we conclude to be the true *cona* ortholog (and used for all analyses). *GJ20698*, which we named *coronetta* (*conta*), appears to be a distant paralog, and, given its presence in the outgroup, emerged prior to the last common ancestor of *Drosophila* and *Scaptodrososphila*. Unlike *cona*, *conta* sequence is conserved (Figure 4A), found in the same syntenic region (in the intron of the gene *teiresias;* Supplementary figure 12), and highly expressed in the testes but not ovaries (Figure 4B). Structural prediction of *CONTA* reveals distinctly shorter coiled structures (Figure 4C). Given the absence of *conta*’s ortholog in *sophophora* – not even in the same syntenic location (Supplementary figure 10) – it has likely been lost, further underscoring the propensity for SC paralogs to participate in germline function that may be evolutionarily fleeting.

**Figure 4.**
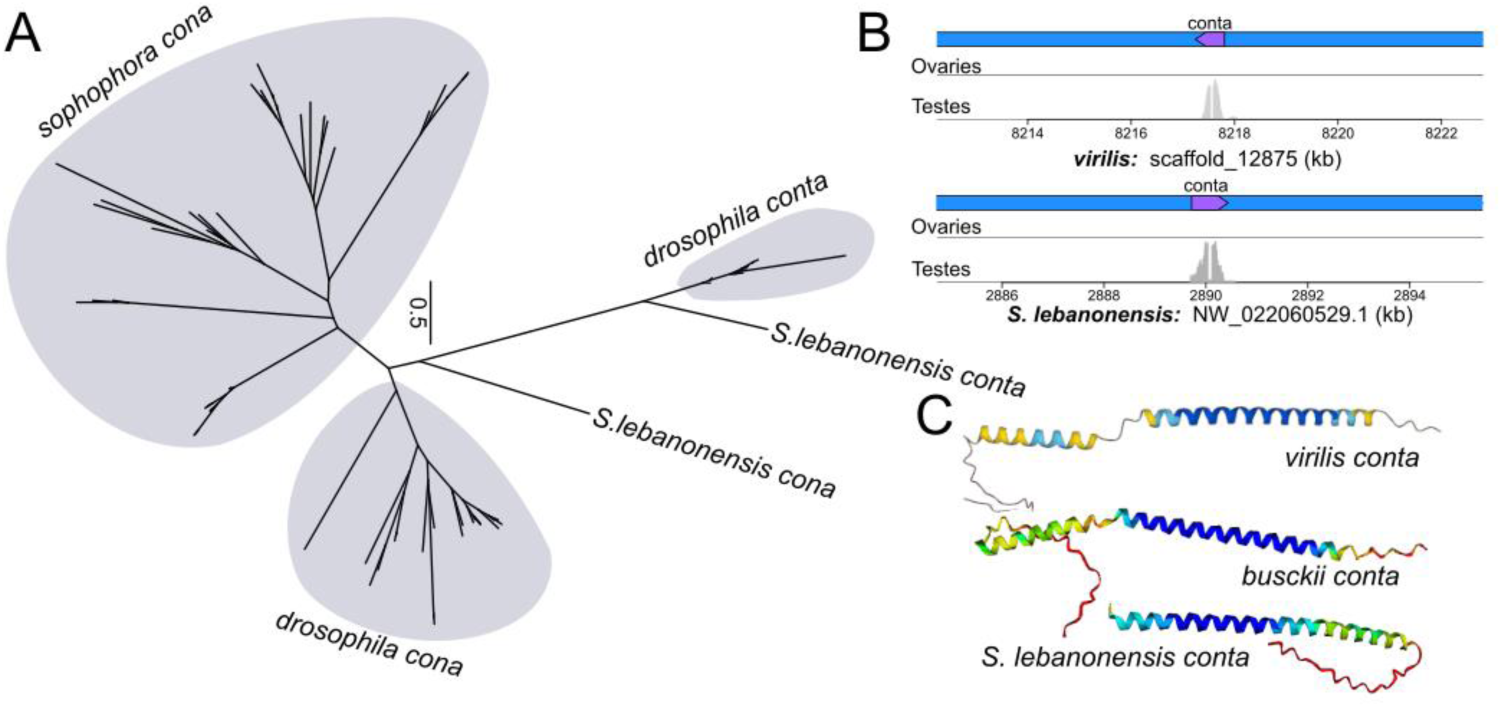
Expression and structure of *conta*, an ancient duplicate of *cona*. A. Unrooted protein tree of *cona* and its old duplicate *conta*; major lineages are labeled. B. Annotation and germline expression of *conta* (purple) in *D. virilis* and the outgroup *S. lebanonensis*. C. Alphafold prediction of conta in representative species.

### SC proteins are evolving under recurrent positive selection and accelerated by repeated duplications

Poor sequence homology can result from relaxed constraint due to reduced negative selection or adaptive protein evolution due to positive selection. Previously, Kursel et al. reported that the elevated rate of SC protein evolution in *Caenohabditis* reflect relaxed sequence constraint while the coiled-coil domains and protein lengths are both highly conserved [25]. However, Hemmer and Blumenstiel identified both elevated rates of protein evolution and signatures of positive selection for *Drosophila* SC genes [12]. We reassessed the rates of protein evolution by estimating the branch-specific ratio of nonsynonymous and synonymous rate of protein evolution, represented by omega. Values approaching 0 indicate negative selection while values close to or greater than 1 indicate relaxed constraint and positive selection, respectively [44]. Notedly, gene-wide omega values, which are predominantly negative, are typically the composite of several modes of evolution as different residues and/or domains of the protein can be under different forms and levels of selection [45,46]. Our curated, species dense SC orthologs and paralogs enabled not only branch-specific, gene-wide estimates (Figure 5A-B and Supplementary figure 12), but also detection of significant positive selection occurring only at portions of the protein coding sequence with the Hyphy package [47].

**Figure 5.**
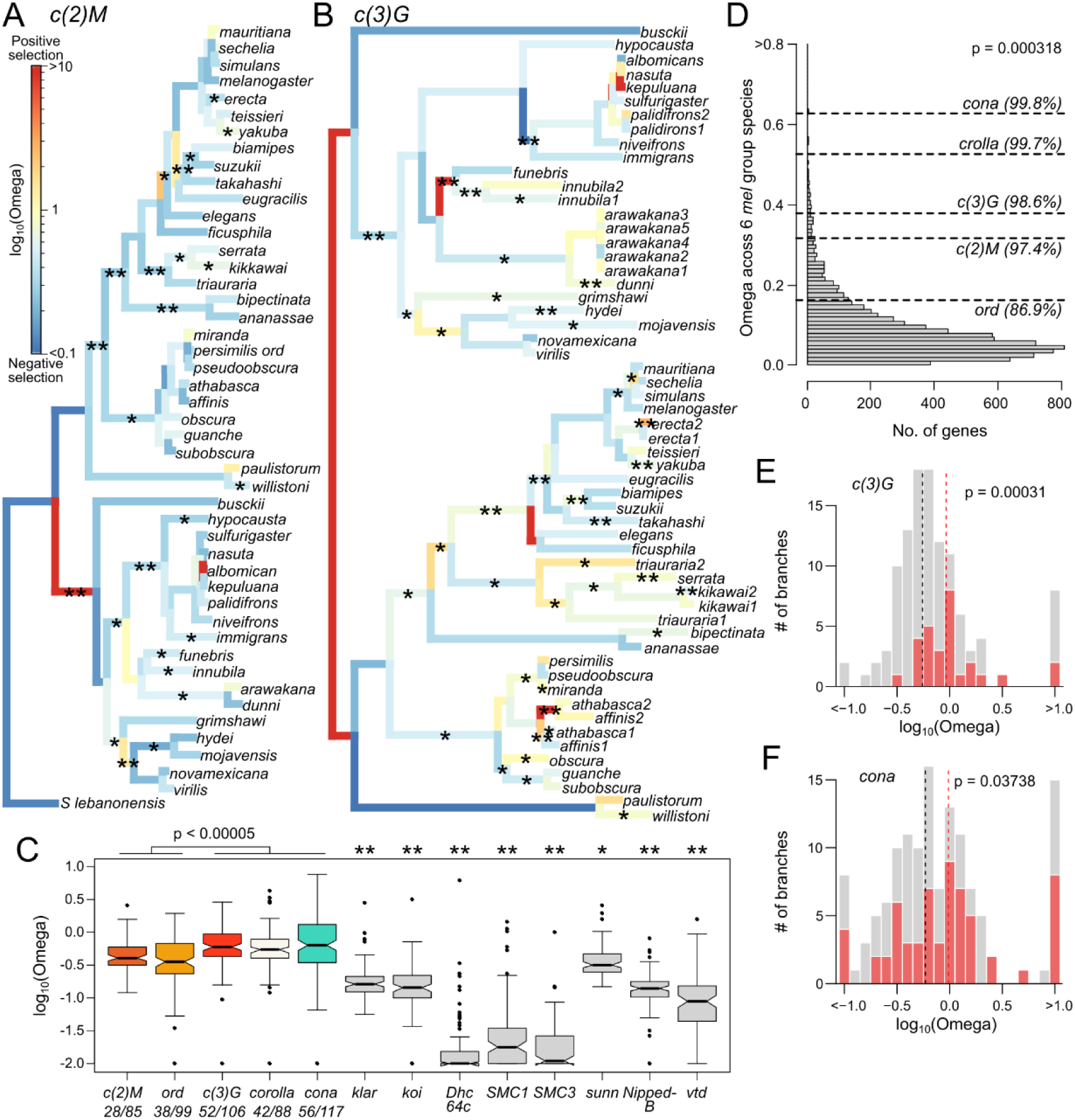
Rate of protein evolution and signatures of positive selection of SC components. A-B. Along the gene trees, branches are colored by their branch-specific rates of protein evolution (omega) with warmer colors representing higher omega. Branches inferred to have significant positive selection in part of the protein are labeled with asterisks (* = p < 0.05 and ** = p < 0.001). See supplementary figure 12 for the remaining SC genes. C. Distribution of branch-specific omega values for the different SC components and genes required for pairing during early prophase I. P-values are from pairwise Wilcoxon’s rank sum tests comparing between the lateral and central region genes. For non-SC genes, ** indicates significant differences (p < 0.005) when compared to all SC genes; * indicates significance comparisons except for *ord*. Ratios below SC genes tallies the number of branches showing omega > 1 or significance over total number of branches. D. *D_N_/D_S_* of the SC genes compared to the genome-wide distribution estimated of by PAML. E, F. Comparison of the distribution of omega values on branches following duplications (red) versus unduplicated branches (gray). Vertical dotted lines mark the median omega values for duplicated (red) and unduplicated branches (black). P-values were inferred from 1-tailed Wilcoxon’s rank sum test.

For the more conserved, lateral elements, despite the better preserved sequence homology and predominantly negative gene-wide omega (Figure 5A and Supplementary figure 13), multiple branches still show signatures of positive selection; 28 out of 85 and 38 out of 99 branches display either positive gene-wide omega or significant site-specific selection for *c(2)M* and *ord*, respectively. For the poorly conserved central elements (Figure 5B & Supplementary figure 12), not only do they have significantly higher omega than the lateral elements (p < 0.00005, pairwise Wilcoxon’s rank sum tests; Figure 5C) indicative of higher rate of protein evolution, over 40% of the branches show either positive omega or significant signatures of positive selection at parts of the protein. Note, we suspect the omega values for *c(3)G*, *corolla*, and *cona* may be underestimated due to poor amino acid alignments across much of the protein. Collectively, the SC shows significantly elevated rates of protein evolution compared to the other genes involved in meiotic pairing; *sunn* is the only exception (Figure 5C) with omega values similar to *ord*, but nonetheless significantly lower than the other SC genes. We further used PAML to infer the rate of protein evolution within the three well represented species groups, and we, again, consistently find evidence of positive selection. Restricting to a subset of 6 *melanogaster* group species with decent alignments, we compared the omega values of SC genes to the genome-wide distribution and found that the SC genes fall between 86.9 and 99.8 percentiles, and are significantly overrepresented with elevated values despite only 5 genes (Figure 5D; p = 0.000318 Wilcoxon’s Rank Sum Test). Altogether these results demonstrate that all components of the SC have a history of recurrent adaptive evolution with the central region genes under frequent and repeated positive selection.

Copy number expansions can allow genes to diversify leading to new functions or subdivision of existing functions among the paralogs. Both of these scenarios are associated with elevated omega, either from relaxed functional constraint or positive selection for novel function. To test whether the recurrent duplications of SC components lead to elevated rates following such functional diversification, we examined branches after duplications for *c(3)G* and *cona.* We found significantly elevated rates of protein evolution on such branches with median omega of 0.925 and 0.914, respectively, both significantly higher than single copy branches (Figure 5E and F; p = 0.000031 and 0.03788, respectively, one-tailed Wilcoxon’s rank sum test). In particular, the *c(3)G2* paralogs with high testes expression in *D. triauraria*, *athabasca*, and *affinis* (Figure 3C-E) all show clear signatures of adaptive evolution post duplication (Figure 5B). Although the terminal branches of the latter two species appear neutrally evolving, the parent branch displays highly elevated and significant omega, indicating strong positive selection in the common ancestor post duplication. Notably, the original copy, *c(3)G1,* in these cases also display signatures of positive selection suggesting that the duplications were followed by functional diversification to both. Moreover, other *c(3)G* duplicates, even the recent ones, show signatures of adaptive evolution including those in *D. erecta* and *kikkawai*. Similar for the *cona* duplicates, nearly all the branches of *cona2* in the serrata group show signatures of positive selection, which accounts for the overall longer branch lengths than those of *cona1* within the same group (Supplementary figure 12). These repeated signatures of positive selection after duplications strongly argue for adaptive functional diversification, acting to further accelerate the already rapid evolution of the SC.

### Poor regulatory conservation of SC genes in both female and male germlines

With the exception of *ord* which maintains sister chromatid cohesion in the male germline [48,49] the *Drosophila* SC proteins are thought to primarily function in the ovaries and dispensable in the testes because males are achiasmic and mutant males do not show obvious meiotic or fertility defects [50–53]. However, a recent elegant study of the male germline by Rubin et al. found that the progression of pre-meiotic chromosome pairing is slower in *cona* and *c(3)G* mutants, revealing putative male germline function despite the absence of SC assembly [54]. Inspired by these results, we examined the expression of SC genes in ovaries and testes RNA-seq datasets from a subset of 38 species (Figure 6A), 14 of which we generated and 26 curated from publicly available datasets (Supplementary table 3). Other than *ord* which is testes-biased across all species, the expression of SC genes – even single copy ones – is highly unstable contrary to the naive expectation of high ovary and low testes expression. testes-biased expression appears to be the norm rather than the exception as SC components are more highly expressed in the testes in over 70% of the species. We further examined available testes single cell RNA-seq data of *D. miranda* [55], a species with high testes expression of *c(3)G*, and find expression concentrated in the pre-meiotic cell types such as germline stem cells and spermatogonia (Supplementary Figure 14), reminiscent of their reported pre-meiotic activity in *D. melanogaster*. Thus, even though they are primarily known for their role in female meiosis, SC genes can also be highly active in testes, arguing for critical male germline function. This is further supported by the heavily testes-biased expression of *corolla*, *c(2)M*, and *ord*, in the outgroup *S. lebanonensis*, suggesting that testes function is ancestral.

**Figure 6.**
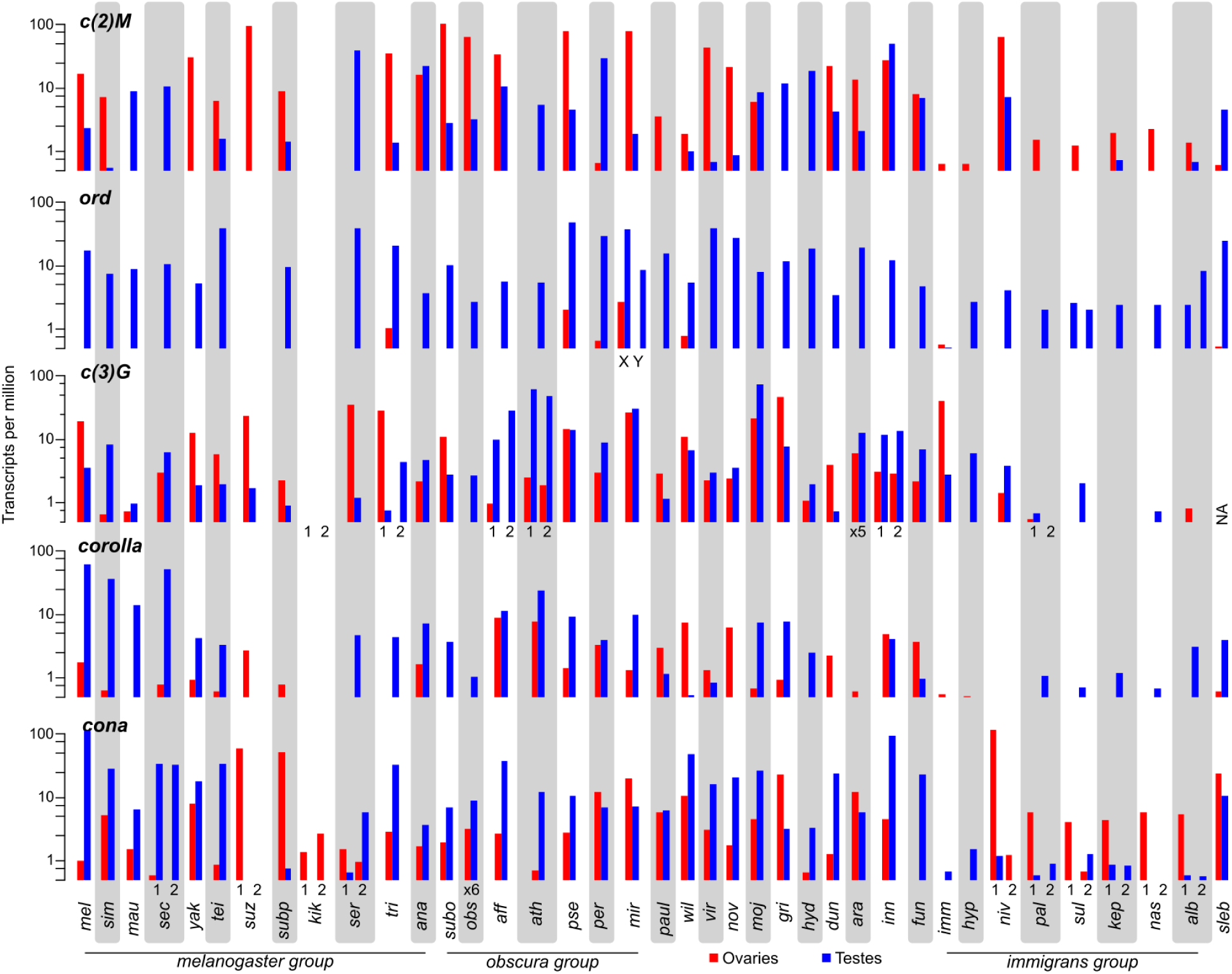
Variable expression of SC components in the testes and ovaries. Ovaries (red) and testes (blue) expression (transcript per million in log scale) of SC genes across 38 species. Tandem duplicates with similar expression are collapsed into one. Other duplicates are labeled. Gray and white vertical bars differentiate between species.

The striking lability of germline SC expression is particularly evident from several closely related species pairs whereby expression rapidly switches between testes- and ovaries-bias. For instance, *c(2)M* is ovaries-biased in *D. simulans* but testes-biased in the sister species *D. sechelia and mauritania* (Figure 6). Similar rapid expression change of *c(2)M* is also observed in *D. pseudoobscura*/*D. persimilis* and *D. athatbasca*/*D. affinis* sister pairs, suggesting such regulatory evolution occurs regularly. As the RNA-seq datasets came from different studies, we tested the possibility that the expression differences resulted from external factors like rearing conditions and examined SC expression from RNA-seq of studies where flies were reared in different environmental conditions. In the datasets examined [56–58], SC expression shows minimal change in different rearing temperature (Supplementary Figure 15). Additionally, drastic expression divergence of the germlines can be observed in RNA-seq datasets we generated of closely related species reared in the same controlled laboratory condition, including those in the *immigrans* group and the sister species *D. arawakana* and *D. dunni*. Therefore, highly variable expression of the SC across the genus likely reflects bona-fide regulatory divergence. Other curious patterns include elevated testes but low-to-no ovary expression such as *c(3)G* in *D. obscura* and *D. hypocausta*. Most puzzling, there are multiple lineages where components of the SC show little-to-no expression in both gonads, such as *D. kikkawai* with little gonad expression of all components. The most extensive regulatory stability appears to be the *nasuta* subgroup where the expression of SC components are consistently low across the species, especially for *c(3)G* and *corolla*. These drastic differences between species cannot be simply driven by differential tissue contribution in the dissections, as the expression of the SC components are poorly correlated, with many instances of high expressions of one component and low expression of others within the same sample. While it is tempting to infer function outside of the germline based on the absence of expression, even *D. melanogaster* ovaries show low expression of some of the SC components like *ord* and *cona* arguing that normal SC function may not require large transcript counts. Nevertheless, these expression patterns reveal that despite the essential roles in regulating crossovers, germline transcriptional regulation of many SC genes is highly labile and constantly evolving, echoing their protein and copy number evolution.

## DISCUSSION

### Trials and tribulations of gene ortholog and CDS searches from genome releases

The species-dense examination of SC evolution was made possible by the large amounts of *Drosophila* genomes that have been recently published. One of the promises of the ever-increasing number of genomes is to enable deep and broad investigation of the molecular evolution of genes and pathways. Naturally, analyses of genic evolution typically require alignments of full length CDSs, which are distilled from genome annotations. Since good gene structure inference requires additional data such as RNA-seq, only ∼1/3 of the available *Drosophila* genomes have available annotations (including those we generated here), which were the focus of this study. Notably, recent studies were able to take advantage of more genomes by focusing on small genes such as protamines [59], sex peptide, and sex peptide receptor [60], which either have no intron or have simple gene structure, enabling protein construction without transcript information from RNA-seq. Even when we relied on annotations with RNA-seq support to identify gene orthologs from annotated genomes using reciprocal best blast hits, we were flummoxed by the regularity of annotation errors (Supplementary table 3). Rapidly evolving genes may be particularly prone to such errors due to obscured sequence homology and, in the case of the SC genes, propensity to duplicate. Without careful manual curation, cross referencing with homology and RNA-seq data, annotations errors taken at face value would be falsely construed as evolutionary changes. Below, we discuss some of the pitfalls of identifying orthologs and paralogs from annotated genomes.

We find that NCBI annotations are generally more reliable than our in-house annotations with maker but has its own sets of issues, the biggest of which is the lack of access to the proprietary pipeline. Problems common to all pipelines include improper splice junctions producing truncated ORFs with premature stops and fused genes creating genes chimeras d – tandem duplicates were particularly problematic for this as they cause extensive mapping errors of the RNA-seq reads between neighboring copies. These errors will cause protein sequences with excess length differences and can be detected as long stretches lacking homology between orthologs in multiple sequence alignments. The more insidious issues stem from genome assembly errors most often occurring as indels at homopolymers tracks, an issue common to Pacbio and Nanopore reads [61,62]. While released genomes should have been polished with Illumina reads to reduce such errors [61], we repeatedly identified annotation errors caused by frameshift indels at homopolymer tracks, the worst offender of which is a Refseq genome, released on and annotated by NCBI, rife with such indels (Supplementary Figure 16). In our in-house annotations with maker, such indels can cause the expected frameshifts and truncated ORFs or unexpected splice junctions with no RNA-seq support. Puzzlingly, NCBI annotations add nonsensically short (<5 bp) introns around such frameshifts to maintain the ORF (Supplementary Figure 15). Because this creates near complete proteins but with small numbers of missing internal amino acids, it is only detectable by looking at variant calls in mapped Illumina reads. Careful examination of CDSs and protein sequences extracted from genome assemblies is therefore necessary to avoid erroneously calling species-specific substitutions, indels, nonsynonmous changes, psuedogenes, and splice variants.

### Mechanisms underlying recurrent duplications and rapid evolution of the SC

Our exhaustive survey for SC orthologs resulted in the surprising identification of many duplicates across *Drosophila*. Duplication is an important mechanism to diversify gene function as the resulting paralogs can either evolve novelty or compartmentalize existing functions, with reduced selective constraint on the gene as a single copy. Indeed, we find elevated rates of protein evolution following duplications indicative of both relaxed constraint and adaptive evolution. Many of the duplicates are young and species-specific showing no evidence of expression differences, and therefore unlikely to have diverged in function. For some of the older *c(3)G* and *cona* paralogs, we observed repeated and independent acquisition of distinct activities surprisingly in the male germline, such as testes-specific expressions and incorporation into lncRNA production. In the case of the tandem duplication of c(3)G in the ancestor of *D. affinis* and *D. athabasca* (Figure 3D-E), the two copies show clear evidence of positive selection driving diversified function evidenced by important structural differences (i.e. loss of terminal globular dominas) in the protein and expression in different testes cell types. While divergence in expression domains is consistent with subfunctionalization, the adaptive protein evolution strongly argues functional novelty. Notably, a parallel duplication *of c(3)G* occurred in *D. triauraria*, which also evolved into a testes-expressed paralog lacking the globular domains that interact with lateral elements. This surprising molecular convergence raises the intriguing possibility of a common testes process acting to repeatedly drive the rapid evolution of SC proteins.

For reasons still unclear, the testes appear to be a unique regulatory environment, producing the largest repertoire of lncRNAs [63,64] and *de novo* genes [65–68], both of which can have critical roles in spermatogenesis. Considering our unexpected finding of frequent testes expression, we speculate that testes activity of SC genes – even single copy ones – may provide the opportunities to generate paralogs that can diversify. This path to functional novelty or diversification in the testes may be further facilitated by the elevated retrotransposon activity in spermatocytes [69] which can conceivably provide the necessary machineries for both retro- and tandem duplications. Our observation that *c(3)G* moved into a satellite-block in the lineages leading up to the pseudoobscura subpseices (Supplementary figure 8) lends support to this possibility.

However, despite many paralogs, some of which with likely novel functions, pseudogenization of duplicates also appears common. We have identified multiple instances where extant duplicates in one species have been lost in neighboring lineages (e.g. *c(3)G2* in *D. triauraria*, and *D. innubila*, and *conta*). We also found several examples of remnants of SC duplicates, including truncated copies of *c(3)G* and *corolla*. Such dynamic copy number changes raise a perplexing conundrum: why are *cona* and *c(3)G* prone to produce duplicates under positive selection only for the duplicates to end up as pseudogenes. One clue may come from the exceptional duplication of *ord* in *D. miranda*, where neo-X- and neo-Y-linked gametologs, along with other meiosis-related genes, have massively amplified in tandem [38]. The amplification is hypothesized to be the result of dosage-sensitive sex-ratio meiotic drivers precipitating an arms race for gene copy numbers on sex chromosomes [38,55]. Similar dynamics of repeated copy number evolution is also observed for sex ratio drivers and suppressors that manipulate DNA packaging in X- and Y- bearing sperms [59,70,71]. In such models of meiotic conflicts, temporary/young duplications may act to increase the gene dosage to either induce selfish transmission (such as biased sex ratio) or to act as suppressors of drive that restore fitness reduction associated with non-mendelian transmission. Once the conflict is resolved, drivers and suppressors may pseudogenize and degenerate as they no longer impart fitness benefits.

Since the intricate orchestration of chromosome movement is necessary for recombination and faithful disjunction, we find the regulatory variability of SC genes in the ovaries to be perplexing, particularly in species where SC genes have little-to-no ovary expression. The repeated relocation to of *c(3)G*, *cona*, and *corolla* to different chromosomal regions may confer some degree of regulatory changes, but even c(2)M, which has remained in the same syntenic location, show variable expression. It is tempting to speculate that the low ovary expression in multiples species correspond to repeated loss of meiotic function, but we find other possibilities more likely. If SC proteins have long half-lives, minimal transcript production may be sufficient to support robust SC assembly. However, this possibility cannot address why species evolved to have drastically different expression profiles. While we examined available germline expression datasets under different environmental conditions and found minimal changes in SC expression, we cannot fully rule out extrinsic factors driving to the expression lability of SC genes. Indeed, recombination rate is sensitive to environmental conditions and life history such as nutrition [72,73], temperature [2,74,75] and stresses [76], and age [77,78], and species can differ in their physiological responses that subsequently regulate SC activity. Such mechanisms can be beneficial in ensuring optimal recombination rates to modulate the amount of genotype diversity in the offspring [76] or proper progression of meiosis in suboptimal cellular conditions like extreme temperatures [9]. However, we note that the species we generated germline RNA-seq for were raised under common laboratory conditions and were at standard, reproductively active age, we still observed drastic differences in SC regulation likely reflecting true regulatory divergence in both the male and female germlines.

Our analyses of the protein coding evolution demonstrate that SC genes have an extensive history of recurrent adaptation with the central region genes being frequent targets of positive selection. This appears unique to SC genes compared to others necessary for chromosomal progression during early prophase which are all conserved, other than *sunn*. While the elevated rates of evolution are partly driven by paralogs diversifying in germline function, orthologs without duplicates also show signatures of positive selection across the gene trees. This contrasts from the SC genes in *Caenorhabditis,* the protein sequence of which are evolving neutrally [25]. Further, while *C(3)G* appears structurally conserved, the lengths of the proteins are far more variable with a coefficient of length variation 5 times higher than that of worms (0.17 vs. ∼0.03 [25]). While this could reflect repeated adaptation in *Drosophila* female meiosis and meiotic recombination, our findings that SC genes frequently function in testes where it is also ancestrally highly expressed compel us to consider additional avenues under recurrent positive selection, especially since spermatogenesis is fruitful grounds for meiotic conflicts, sexual selection, and molecular innovations. Pleiotropy tends to increase molecular constraint [79]. However, given the sequence tolerance of the SC, dual function of *Drosophila* SC genes in both oogenesis and spermatogensis may instead predicate a unique scenario where positive selection in the latter has little pleiotropic impact on the former. Dissecting the function of SC genes in the *Drosophila* male germline, which is ironically achiasmate, will therefore be critical to understanding the diversity and evolution of meiotic recombination.

## MATERIALS AND METHODS

### High molecular weight DNA extraction and genome assembly

To assemble the genomes of *D. hypocausta* and *D. neivefrons*, we followed the Nanopore long read sequencing pipeline from [33,80]. In short, high molecular weight DNA was extracted using the Qiagen Blood & Cell Culture DNA Midi Kit from ∼100 males of *D. hypocausta* strain 15115-1871.04 from the National Drosophila Species Stock Center and ∼50 females of *D. niveifrons* strain LAE-276 from the Kyorin Drosophila Species Stock Center. DNA strands were hand spooled after precipitation, followed by gentle washing with supplied buffers.

### RNA-seq preparation and analyses

5 pairs of ovaries and testes were dissected from adult females and males and stored in Trizol at -80 degrees, followed by standard RNA extraction. RNA-seq libraries were generated using either the NEBNext RNA Library Prep Kit for Illumina with the Stranded and mRNA isolation Modules or the Illumina Truseq Stranded mRNA Library Prep kit. After quality check with the Fragment Analyzer at QB3-Berkeley, the libraries were sequenced by Novogene. We aligned the reads (both ones we generated and downloaded from SRA) using hisat2 (v2.2.1) [81] on either pair-end or single-end mode to their respective genomes with the –dta flag to allow for downstream transcriptome assembly. After sorting the aligned reads with samtools (v1.5) [82], we used the featureCount (v2.0.3) in the Subread package [83] for read-counting over genes, allowing for non-uniquely mapped reads (-M flag). Read count tables were processed and analyzed in R (v4.2.2) and Rstudio (v2022.12.0). For gene expression analyses, we normalized the read counts across samples by converting them to transcript per million (TPM) [84]. For species where we needed to do de novo gene annotation, we used stringtie v2.1.6 [85] on default for genome-guided transcript assembly.

### Gene annotation and manual curation of gene structures

For species that required gene annotation, we ran three rounds of maker [86]. For evidence-based ab initio gene prediction in the first round, we supplied the transcript assembly from stringtie, de novo repeat index from RepeatModeler2 [87], transcript sequence from closely related-species and protein sequence data from D. melanogaster and D. virilis downloaded from FlyBase. The maker results from round one were used to train the species specific gene model using SNAP [88]. The resulting snap.hmm file was fed back into maker for round 2. We iterated this process again, refining the gene models for a 3rd round of maker.

For malformed or missing annotations, we first visualized the gene structures and RNA-seq reads mapping around them using IGV (v2.16.0) [89]. Additionally, we manually defined the region of the genome showing gene homology by blastn-ing the well-formed ortholog from a closely related species to the genome. The combination of RNA-seq reads mapping the blast-hit boundaries provided evidence to correct erroneous exon-intron injunctions, truncated annotations, chimeric gene structures, and absent annotations. To update the annotations file (.gff file), we used the genome browser GenomeView (v2250) [90] to manually edit or add the gene structures including mRNAs, exons, CDSs. All edited genes have at least full open reading frames, although 5’ and 3’ UTRs are missing. All manually annotated features were marked by the flag “hand” in the gffs. The updated .gff is then exported and sorted using GFF3sort [91], and transcript sequences are retrieved using gffread (0.9.12) [92]. In several instances, we noticed assembly errors leading to malformed genes. One was *ord* in D. nasuta which had a stretch of N’s within the gene body indicating scaffolding points. The other was in D. *neivifrons* where c(3)G was annotated as two fragments. This was due to a deletion of a single nucleotide in the genome causing a shifted reading frame which led to malformed annotations. The deletion was revealed by RNA-seq read mapping, whereby all reads showed a one basepair insertion. We fused the fragmented annotations into one, and corrected the transcript sequence to rectify the erroneous deletion. Lastly, we initially could not identify *corolla* in the primary NCBI genome assembly of *D. funebris* strongly suggesting gene is loss; however, we were subsequently able to identify it in an unplaced repeat-rich contig in a separate assembly. For species with malformed annotations that had a closely related species with well-formed orthologs, we used the program Liftoff [93] to convert the annotation from one genome to the other.

### Homolog search with reciprocal best blast hits of transcripts and/or coding sequences

We first used blastn to identify homologous transcripts between species pairs, and in cases where no clear best hit was identified, we then used tblastn to identify homologous transcript with protein sequences. The absence of protein hits with tblastn is then followed by blasting to the genome, to ensure the absence of an ortholog in the transcript sequences is not merely the result of absent annotation.

To identify orthologs and paralogs using a reciprocal best blast hit strategy, we reciprocally blastn-ed transcript sequences from species pairs using the commands:

blastn -task blastn -query species1.transcripts -db species2.transcripts -outfmt "6 qseqid sseqid pident length qlen slen mismatch gapopen qstart qend sstart send evalue bitscore" -evalue 1

blastn -task blastn -query species2.transcripts -db species1.transcripts -outfmt "6 qseqid sseqid pident length qlen slen mismatch gapopen qstart qend sstart send evalue bitscore" -evalue 1.

For publicly available genomes with annotation files, we generated the transcript sequences using gffread, otherwise we used transcript sequences generated by maker. We then used grep to identify the blast hits and checked whether they are reciprocal best hits of each other. For the *Sophophora* and *Drosophila* sub-genera, we used *D. melanogaster* and *D. virilis* sequences downloaded from flybase as the focal species and blasted them first to their close relatives. When one species yields no blast hit for a gene, we then use other closely related species where the orthologs was successfully identified. If no hits can be identified for a species or a clade, we then repeat the same procedure using tblastn to identify translated protein sequences, as amino acid can be more conserved than nucleotide sequence. If tblastn fails to identify a homologous transcript, we then blastn-ed to the genome sequence. True absences/loss of a gene will yield poor or no blast hits, while missing annotation will result in clear noncontiguous hits with gaps corresponding to introns.

### Microsynteny surrounding homolgs and chromosome placement

We extracted the sequences of the homologs including 50kb up and downstream using bedtools slop and bedtools getfasta. We then pairwise blastn-ed the sequences to each other and filtered out alignments with E-values of < 0.01 or shorter than 100 bp. To infer the extent of homology in the flanking sequences, we calculated the proportion of sequence aligned, excluding the positions of the homolog. Genes are deemed to be in non-syntenic regions if they share < 5% flanking homology. For species without Muller element designation of chromosomes, we assigned Muller elements by blastn-ing to the genome of a closely related species where the Muller elements have been determined.

### Phylogeny construction

We retrieved the CDS for all genes, removed the stop codon, and converted them first to protein sequences using EMBOSS Transeq [94]. We then aligned the protein sequences using three aligners with the commands: prank (v.170427) -protein -showtree [95], mafft (v7.505) --localpair --maxiterate 1000 [96], and muscle (v5.1) [97]. The resulting multi-sequence alignment fasta file were used as the input for iqtree (v1.6.12) [98] with the flags -AA and -bb 2000 for 2000 iterations of ultrafast bootstrapping [99]. These trees were manually rooted with *S. lebanonensis* as the outgroup species in FigTree (v1.4.4) [100], and then Node labels added with phytools (v1.5.1) [101] to the trees to facilitate downstream rate of evolution analyses with Hyphy. We then selected the resulting trees with the best bootstrap support and concordance with species tree. *cona* trees were highly inconsistent across alignment methods with many poorly supported nodes.

### Branch-specific rate of protein evolution and positive selection with HyPhy

We used TranslatorX [102] to align the CDS sequence based on the protein alignments. Providing the CDS alignments and the protein trees, we used the ABSREL module in HyPhy (v2.5.51) [47,103] to infer the branch-specific rate of protein evolution (Omega) and significant signatures of positive selection. We wrote a custom script (github.com/weikevinhc/phyloparse) to parse the HyPhy .json output in R where the trees were reoriented with phytools and visualized with colors representing gene-wide omega values. The nominal p-values were used for significance. In addition, we used PAML [104] for the species-group specific test of recurrent positive selection. For comparisons with the rate of regulatory evolution, we reran HyPhy after removing sequences from species with no RNA-seq data.

### Rate of protein evolution with PAML analyses

From the genus wide alignment of SC genes, we extracted six species, *D. melanogaster, D. simulans, D. sechelia, D. yakuba, D. erecta, and D. ananassae*, the same six *Drosophila* species used to calculate dN/dS previously in Clark et al 2007 [105]. We then constructed maximum-likelihood trees using iqtree using parameters ’-m MFP -nt AUTO -alrt 1000 -bb 1000 -bnni’. We then used the PAML parameter (model = 0 and CodonFreq = 2) to estimate the rate of protein evolution across the six species.

### Protein structure prediction with AlphaFold

Structures of proteins previously annotated in NCBI were retrieved from the AlphaFold Protein Structure Database [41]. For genes we annotated, we used ColabFold (v1.5.2), an implementation of AlphaFold on the Google Colab platform [42] and selected num_recycles 24, producing structure predictions that were visualized in UCSF ChimeraX [106].

## Data availability

All generated raw sequences have been deposited onto the SRA under PRJNA1030345 (to be released upon publication). Relevant intermediate datasets and files have been deposited on dryad currently available to reviewers at: https://datadryad.org/stash/share/V4wnkUUlEi_RkeK1XMF9qREOcynzDvA2QjmvjrrAKgU

