## Supplementary figures for "Diversification and recurrent adaptation of the synaptonemal complex in *Drosophila*"

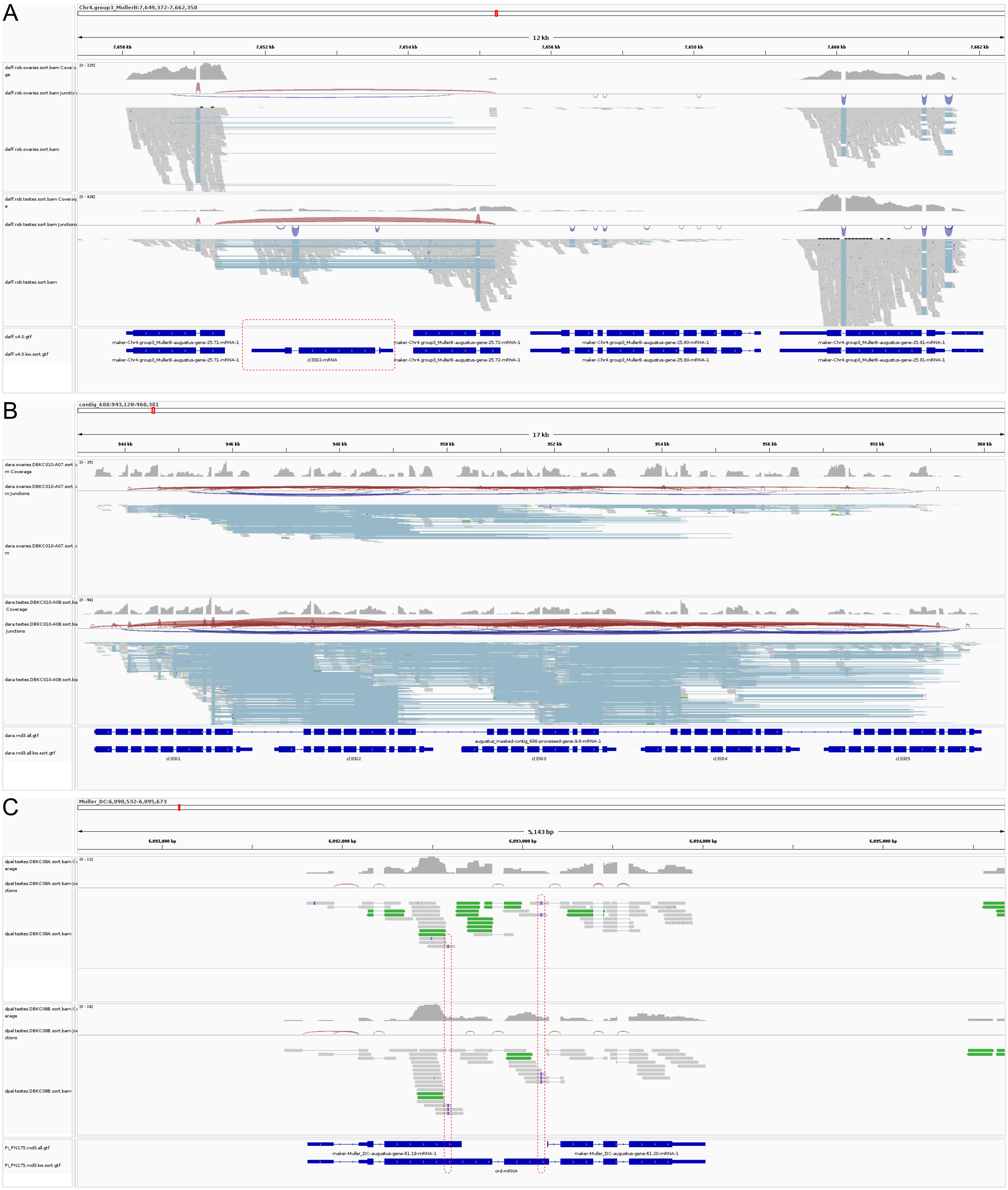


**Supplementary Figure 1:** IGV genome tracks showing examples of common sources of annotaion errors. RNA-seq read depth, splice junctions, and reads are shown. Erroneous and corrected annotations are the two bottom tracks, respectively. A. Absent annotations despite RNA-support (circled by a red box). B. Neighboring genes are fused into a long chimeric gene.  C. Annotation error due to assembly errors that cause frameshifts. RNA-seq reads mapping all show a 1bp insertions (purple marks on the reads and circled by red boxes) indicating that the assembly has misassembled these regions introducing two 1bp deletions. Such errors in the exons destroys the ORF leading to errorneous inferences.

A D. miranda X-linked ord tandem copy regions: MullerC:12,770,000-12,830,000

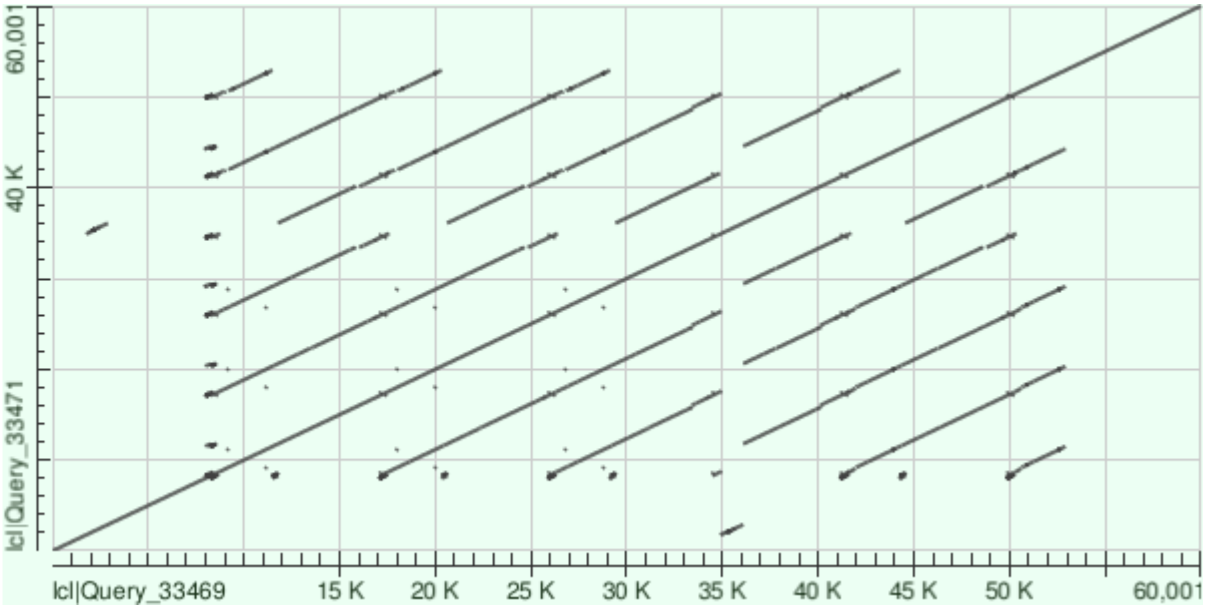


B D. miranda Y-linked ord region: Contig_Y1:27,200,000-27,700,000

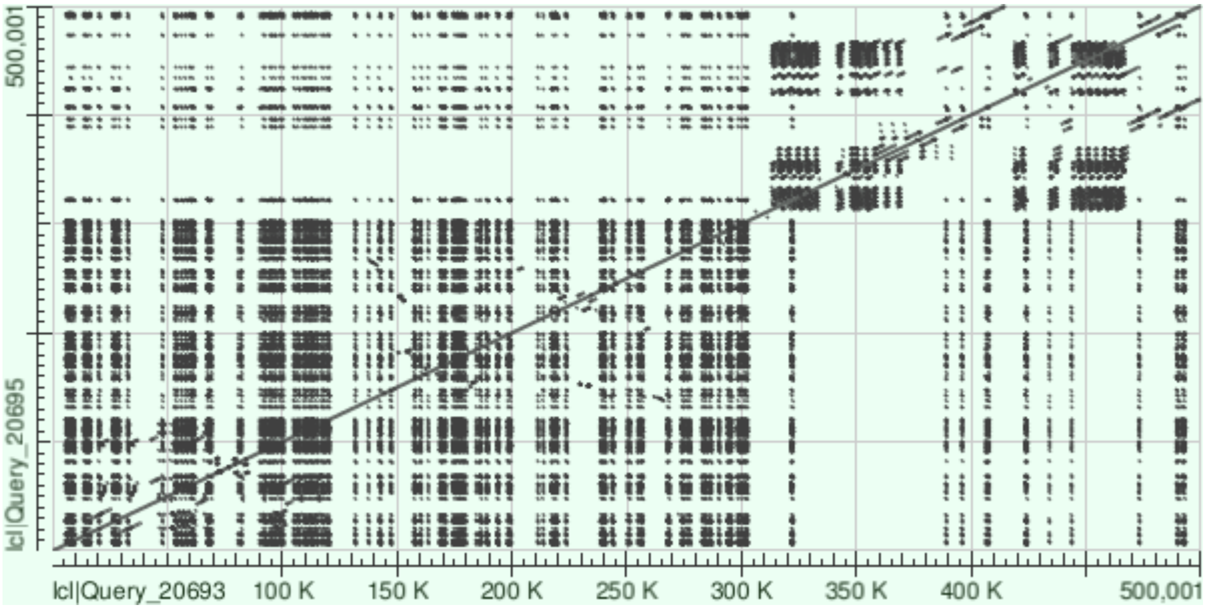


Tandem ord duplications on the neo-X (A) and neo-Y (B). Self-alignments (blastn) for the genomic regions containing ord copies.

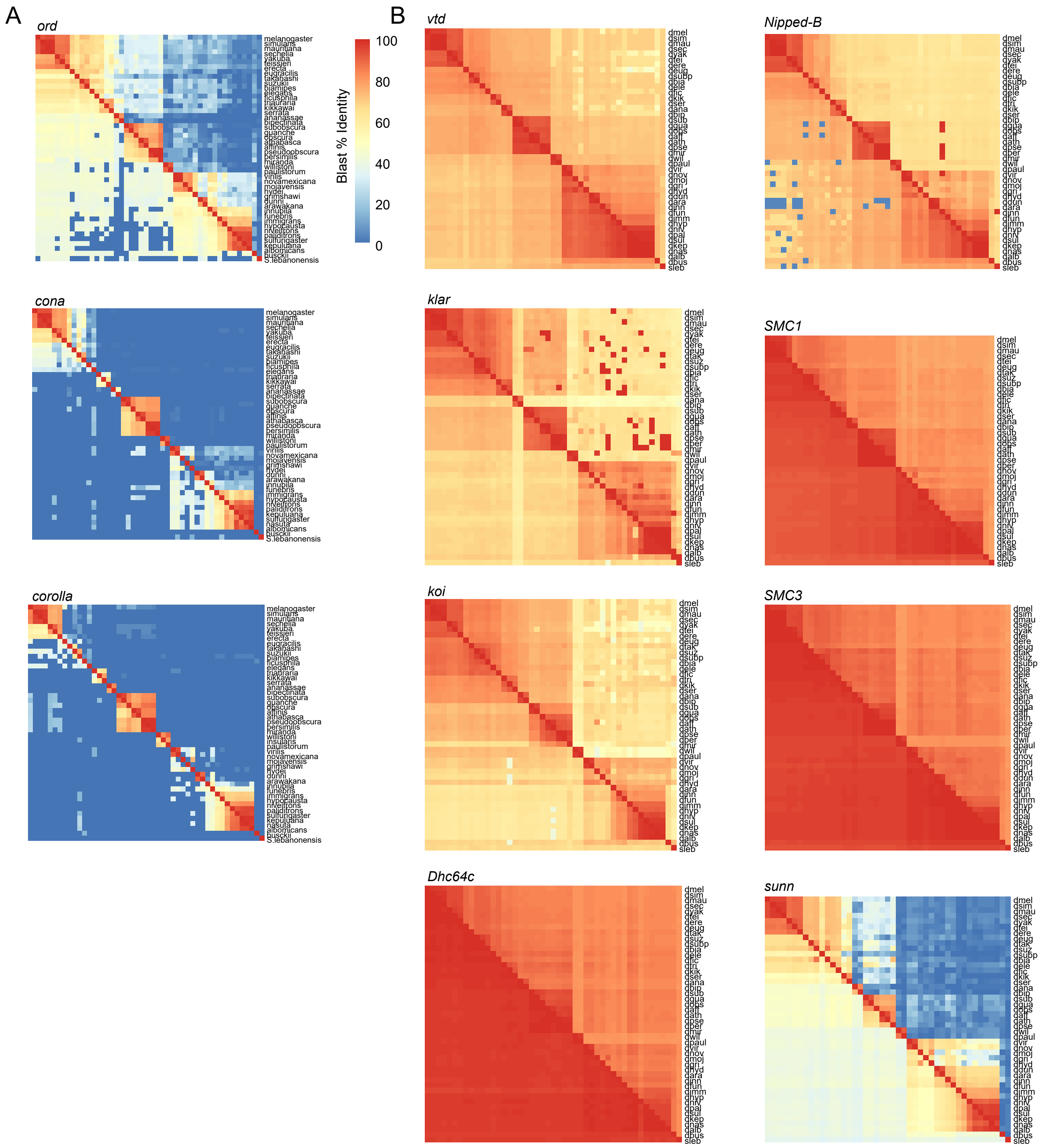


**Supplementary Figure 3:** Sequence similarity of orthologs of (A) SC genes and (B) genes involved in meiotic pairing during female meiosis as measured by blast (same as Figure 1D-F). Pairwise % identity values above and below the diagonal were estimated based on blastn of the CDS and blastp of the AA sequences, respectively.

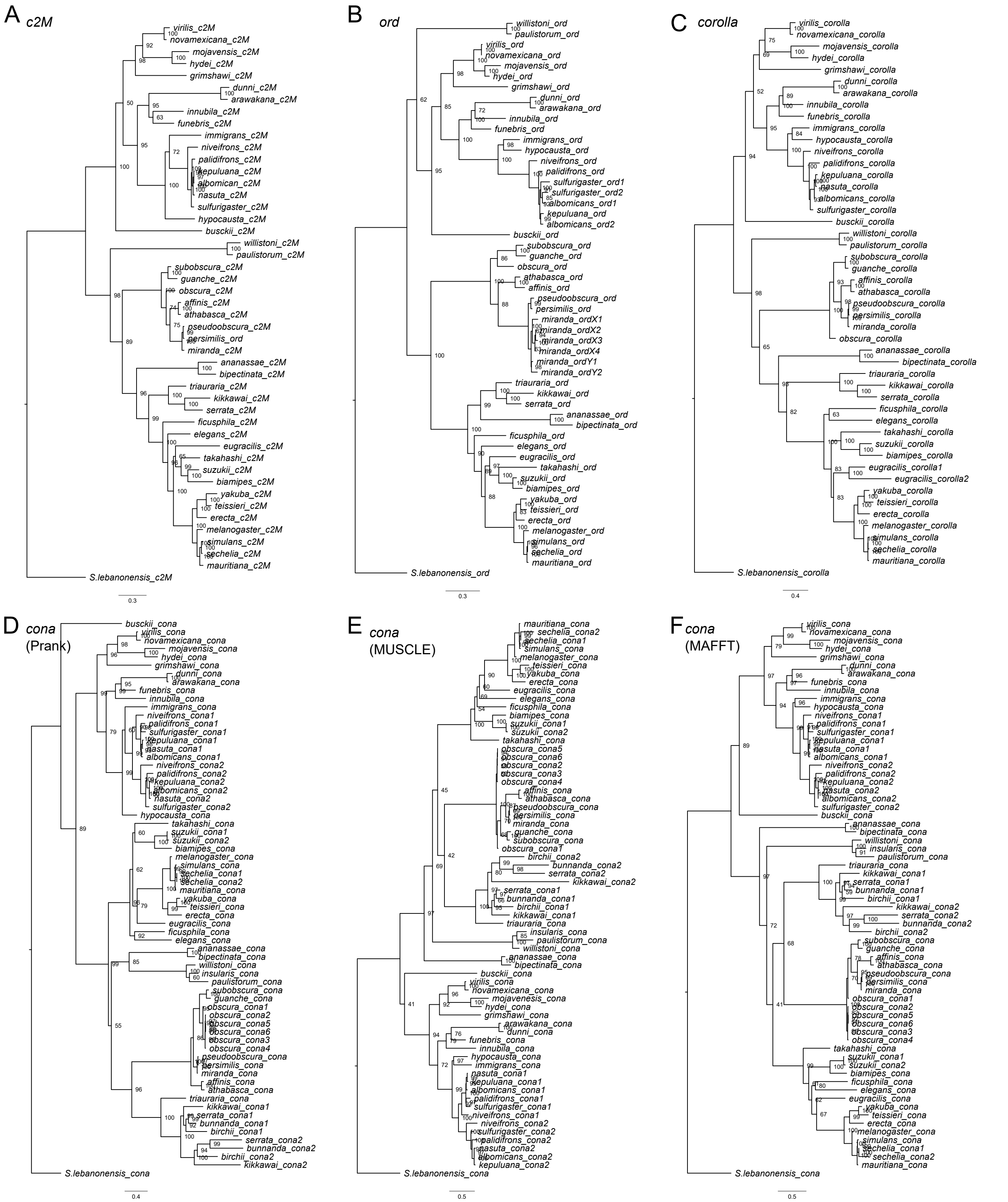


**Supplementary Figure 4:** Gene trees for c(2)M (A), ord (B), corolla (C) and cona (D-F) constructed using the protein alignments. Trees for cona are based on Prank (D), MUSCLE (E), and MAFFT (F) protein alignments.

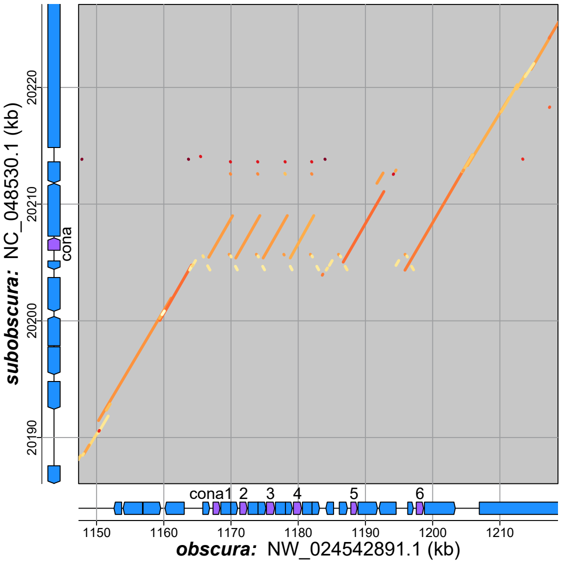


**Supplementary Figure 5:** Six tandem duplicates of cona in

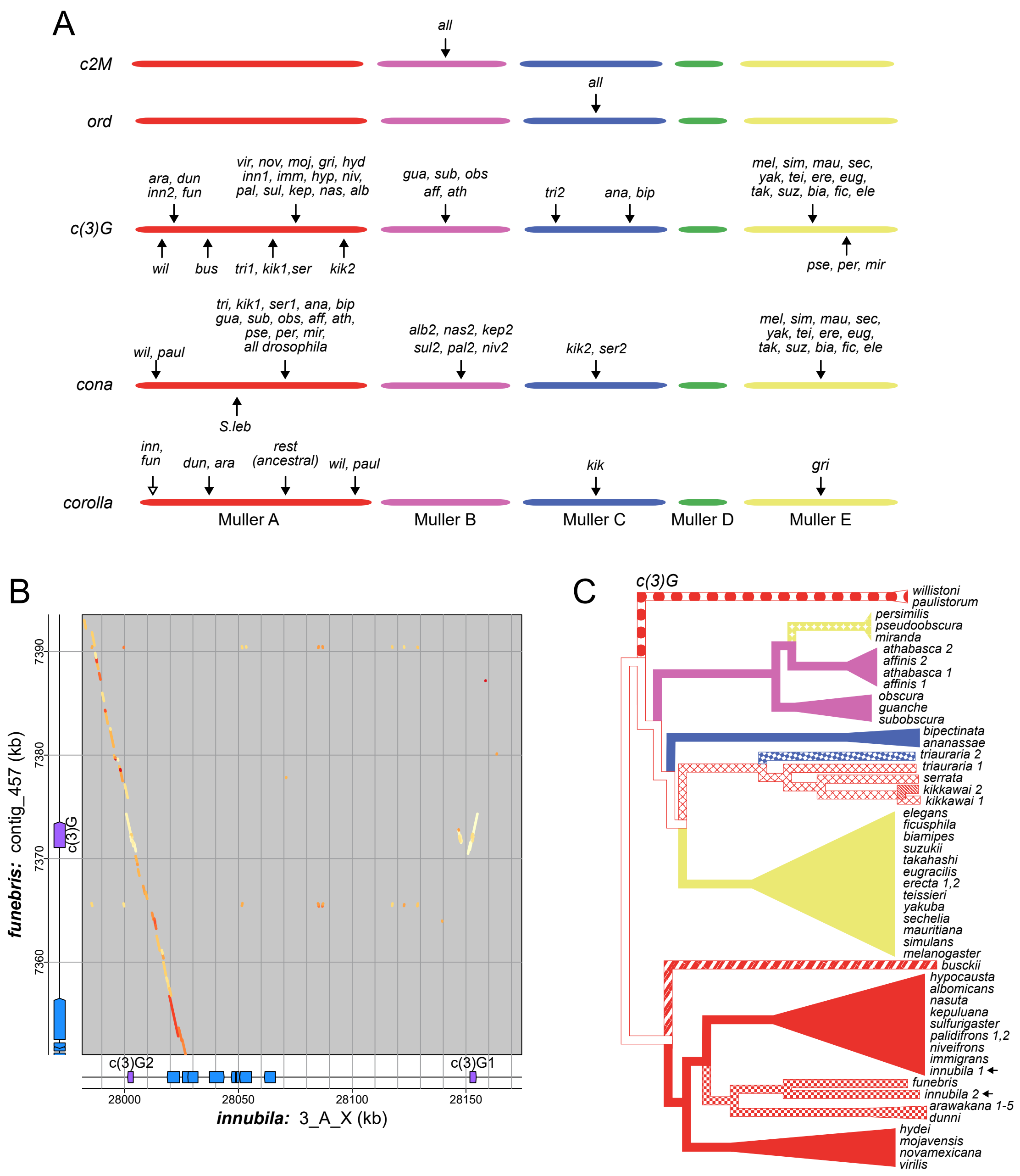


**Supplementary Figure 6:** Muller elements in which SC components are found are labeled. Each arrow indicates a syntenic region where orthologs are found. Muller elements are labeled by different colors and are not drawn to scale. Order of the arrows do not reflect their relative chromosomal locations. Open arrow for corolla indicates insertion into repeat rich pericentromeric regions. B. Dotplot of the genomic region surround c(3)G paralogs in D. innubila compared to D. funebris. Phylogenetic reconstruction of c(3)G position and movements in the genome. Color of branches indicate the Muller elements in which c(3)G resides. Different patterns represent different, non-syntenic locations, on the Muller elements. Arrows point to the D. innubila duplicates which are found in ancestral and derived regions suggesting an old duplication prior to species split with D. funebris.

A

B

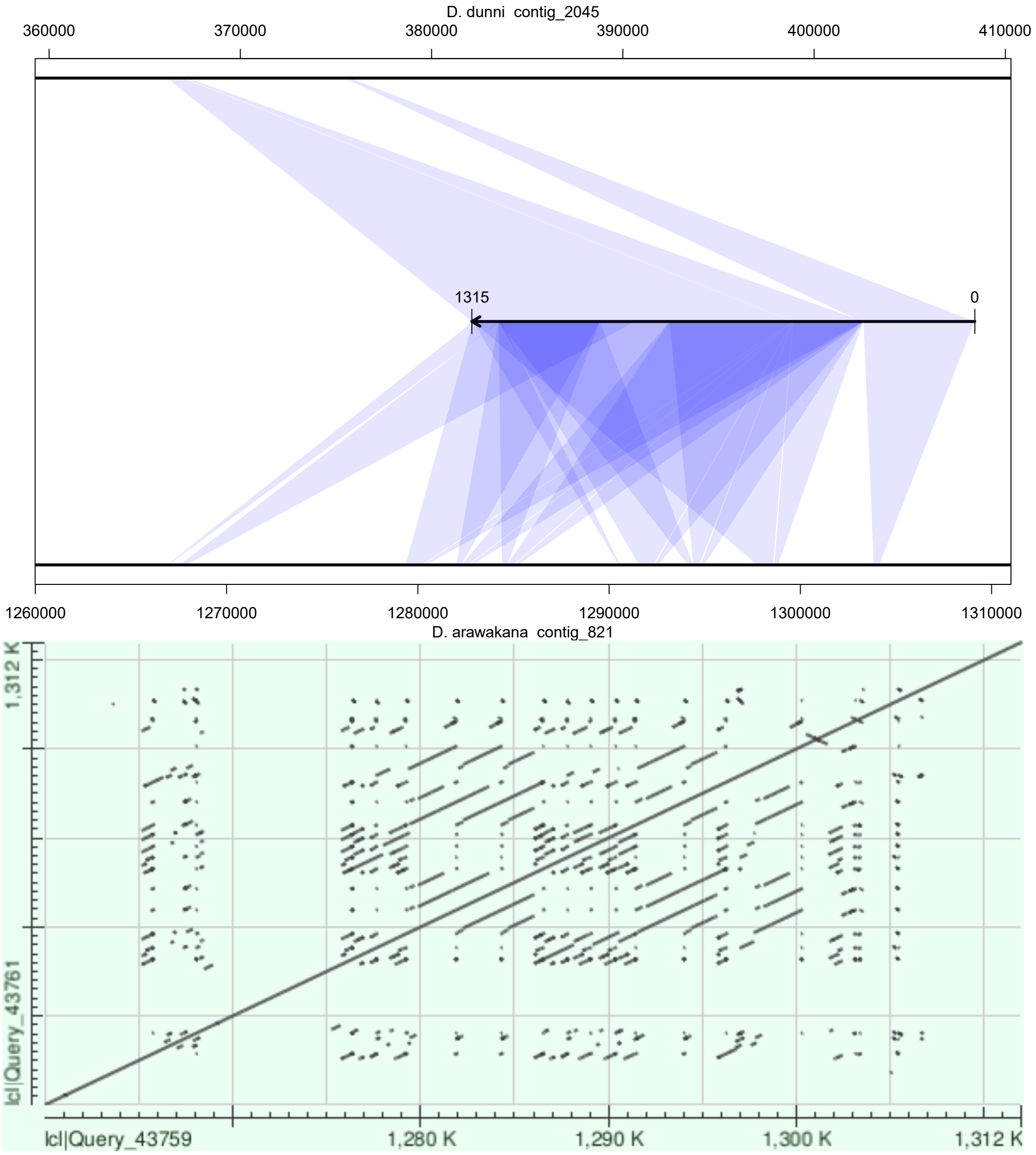


**Supplementary Figure 7:** Truncations of tandem copies of corolla in D. arawakana. A. Alignment of corolla CDS (center track) to the D. dunni (top track) and D.arawakana (bottom track) genomes. B. Self alignment of the genomic region containing corolla revealing complex tandem repeat structures.

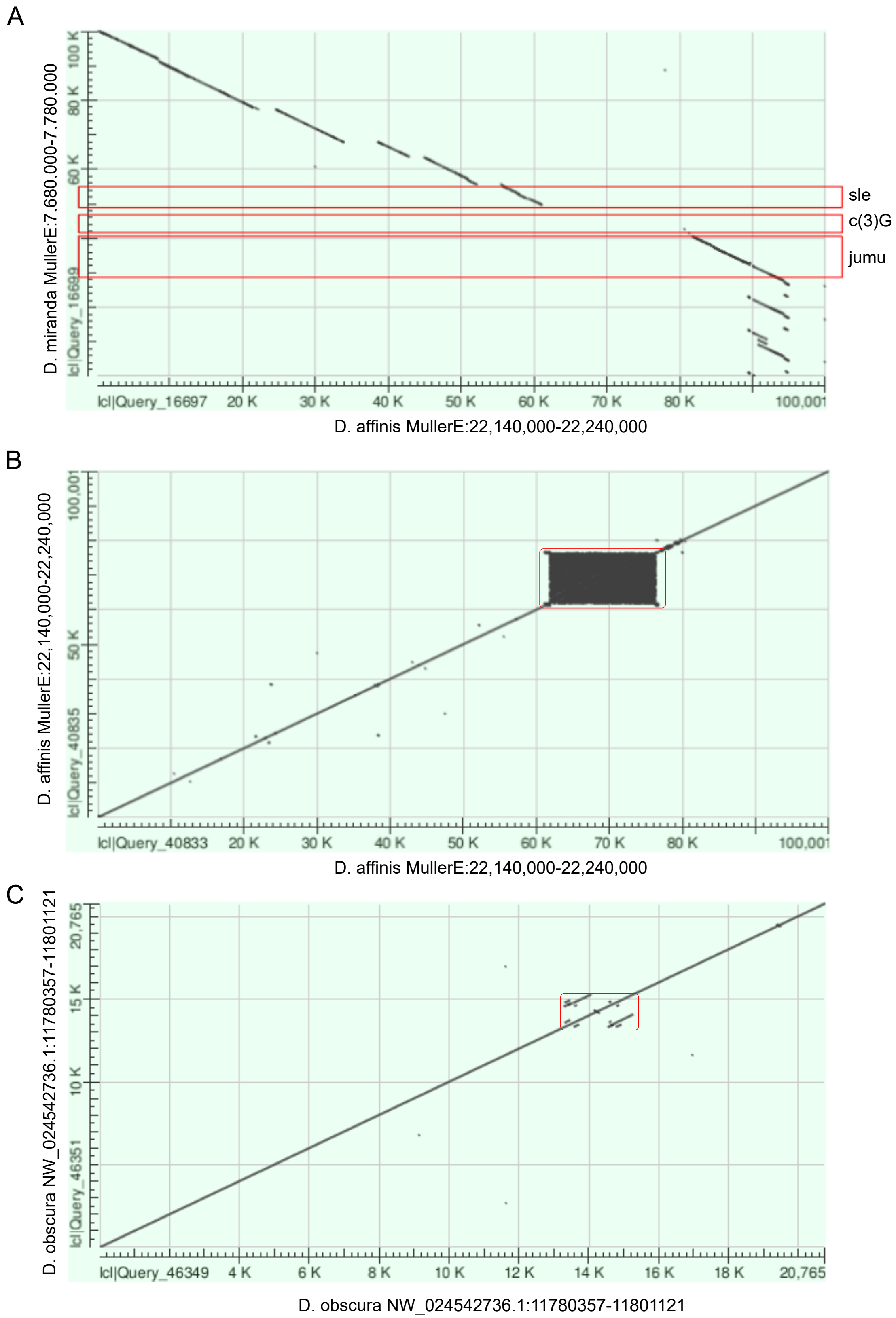


**Supplementary Figure 8:** c(3)G movement in the pseudoobscura subgroup. A. c(3)G moved to Muller E in the pseudoobscura subgroup. Blastn alignment between the c(3)G location in D. miranda to the syntenic region in D. affinis which lacks c(3)G. c(3)G and flanking genes are boxed in red. B. Self alignment of the region c(3)G migrated to show extensive tandem repeat structure in D. affinis. Region where c(3)G inserted into boxed in red. C. Self alignment of the syntenic region in D. obscura.

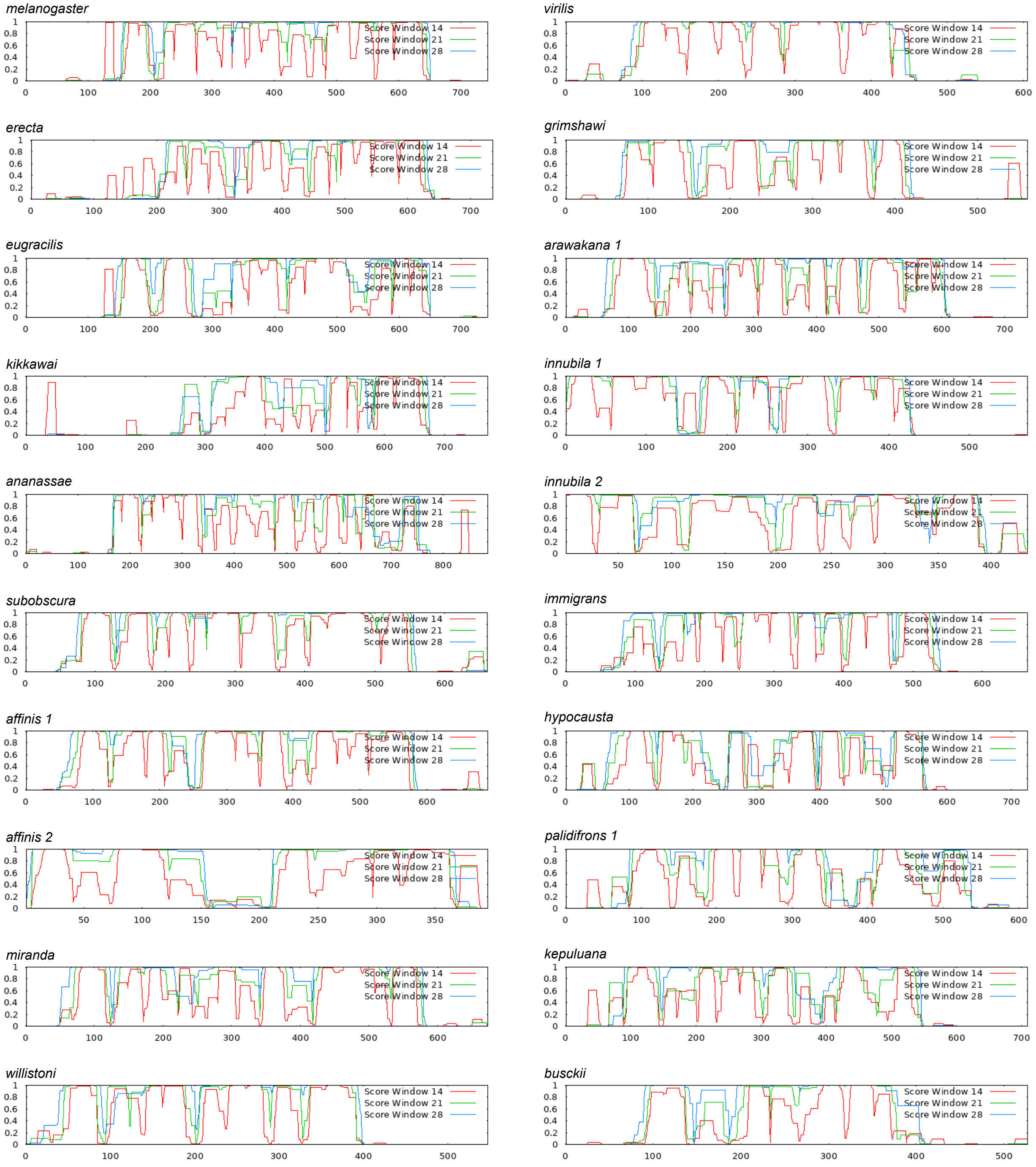


**Supplementary Figure 9:** Coiled-coil domain prediction for representative c(3)G orthologs and paralogs.

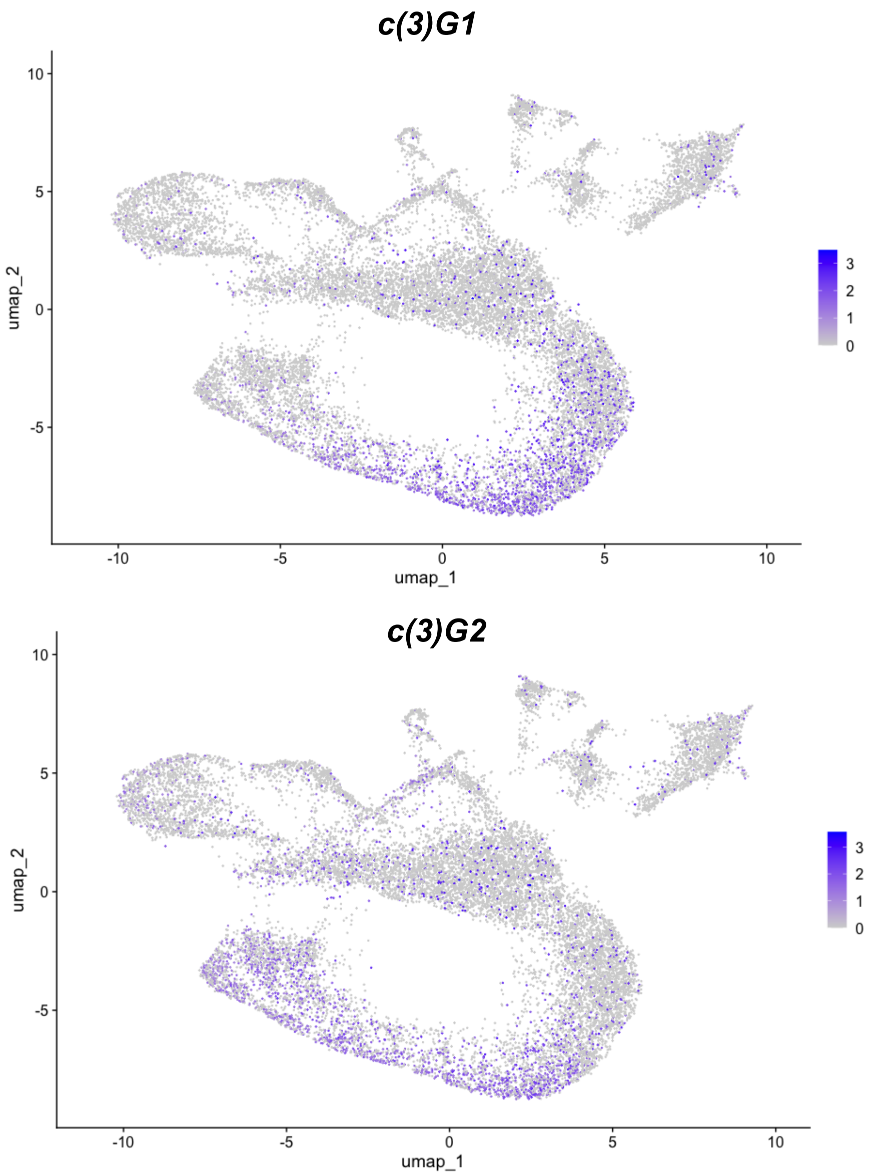


**Supplementary Figure 10:** Expression of c(3)G1 and c(3)G2 in D. affinis single nuclei RNA-seq dataset (Personal Communications with RU).

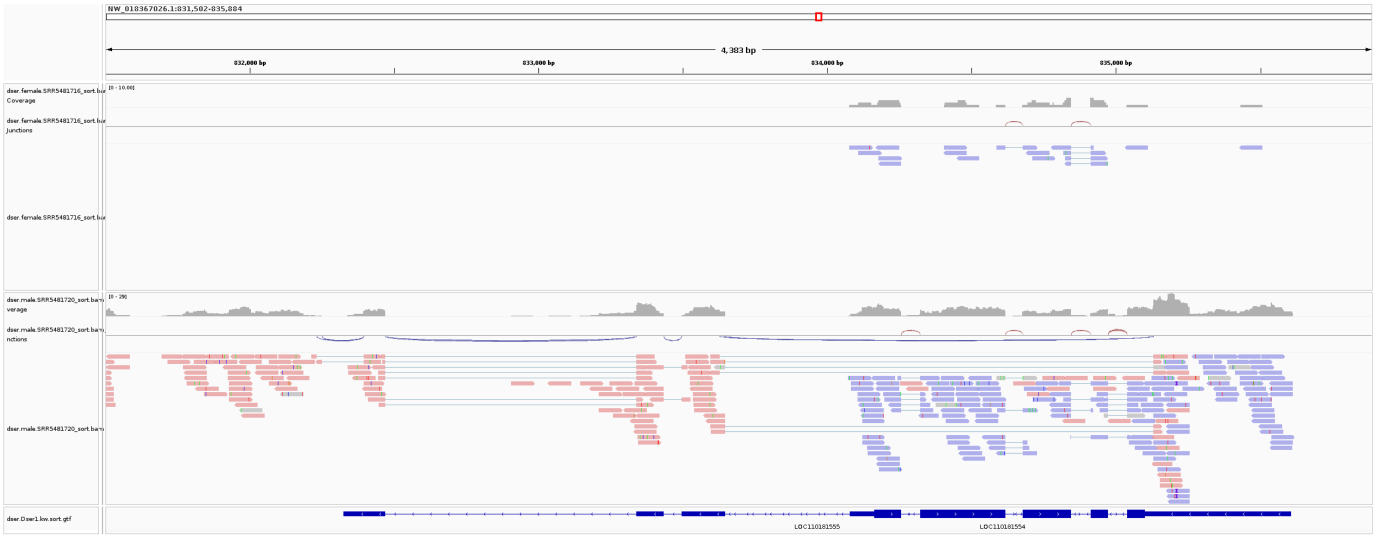


**Supplementary Figure 11:** IGV genome tracks of D. serrata showing expression surrounding cona2 (marked by purple reads) and the anti-sense lncRNA (marked by pink reads). Pink and purple differentiate reads originating from the sense and antisense strands. Note the genome track is in the reverse orientation compared to Figure 3D.

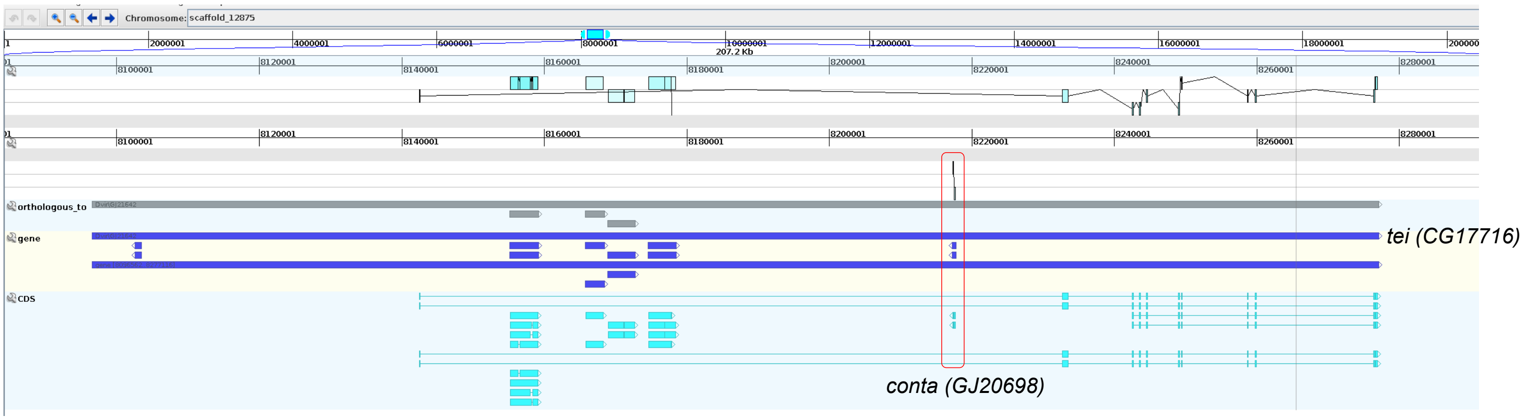


**Supplementary Figure 12:** *conta* location in the *D. virilis* genome. It is embedded in the intron of the gene *teiresias*. Note the lack of ortholog in the gray track.

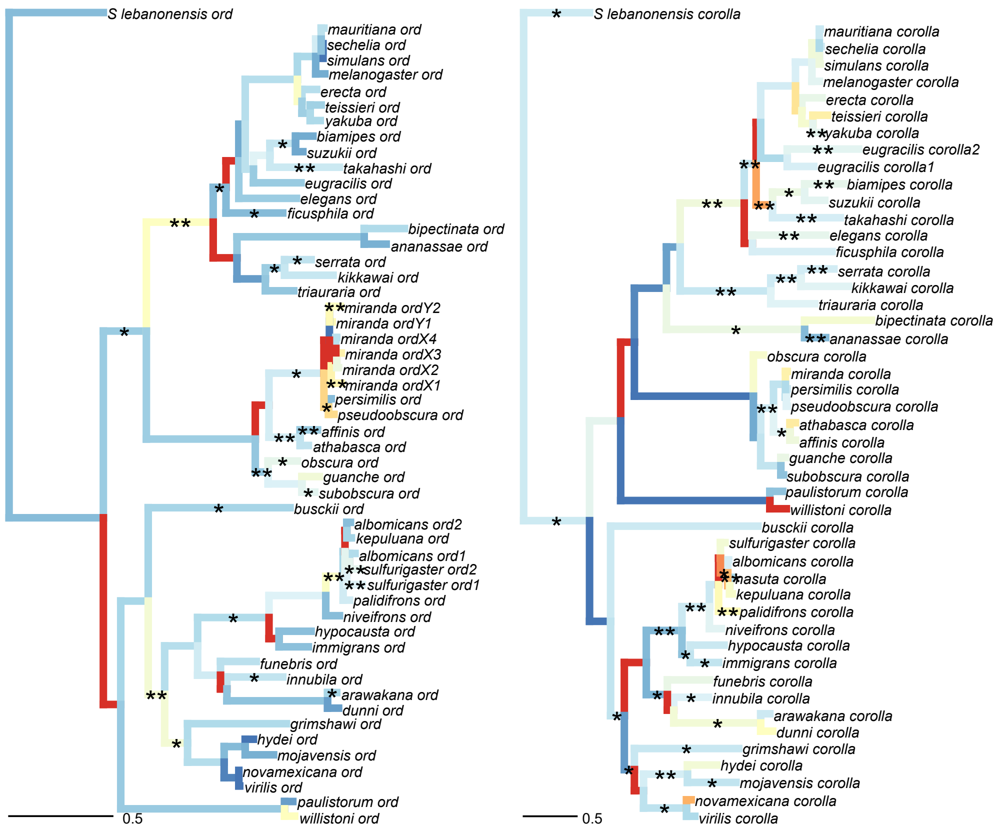


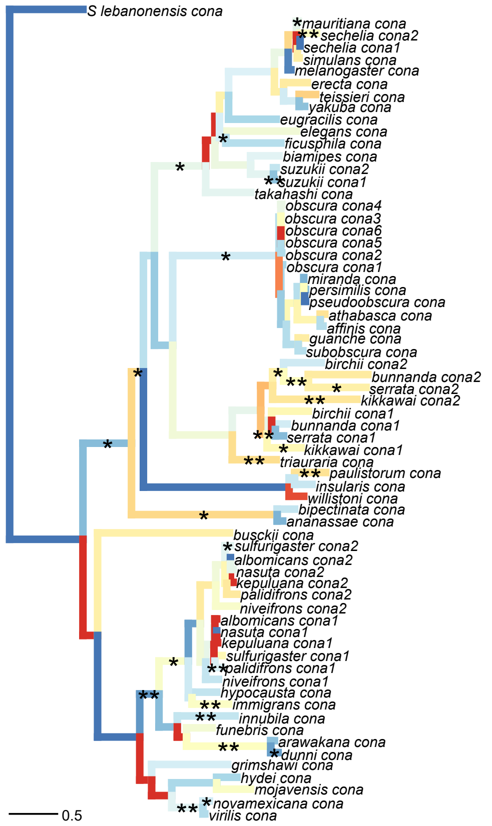


**Supplementary Figure 13:** Branch specific Ka/Ks for ord, corolla, and cona

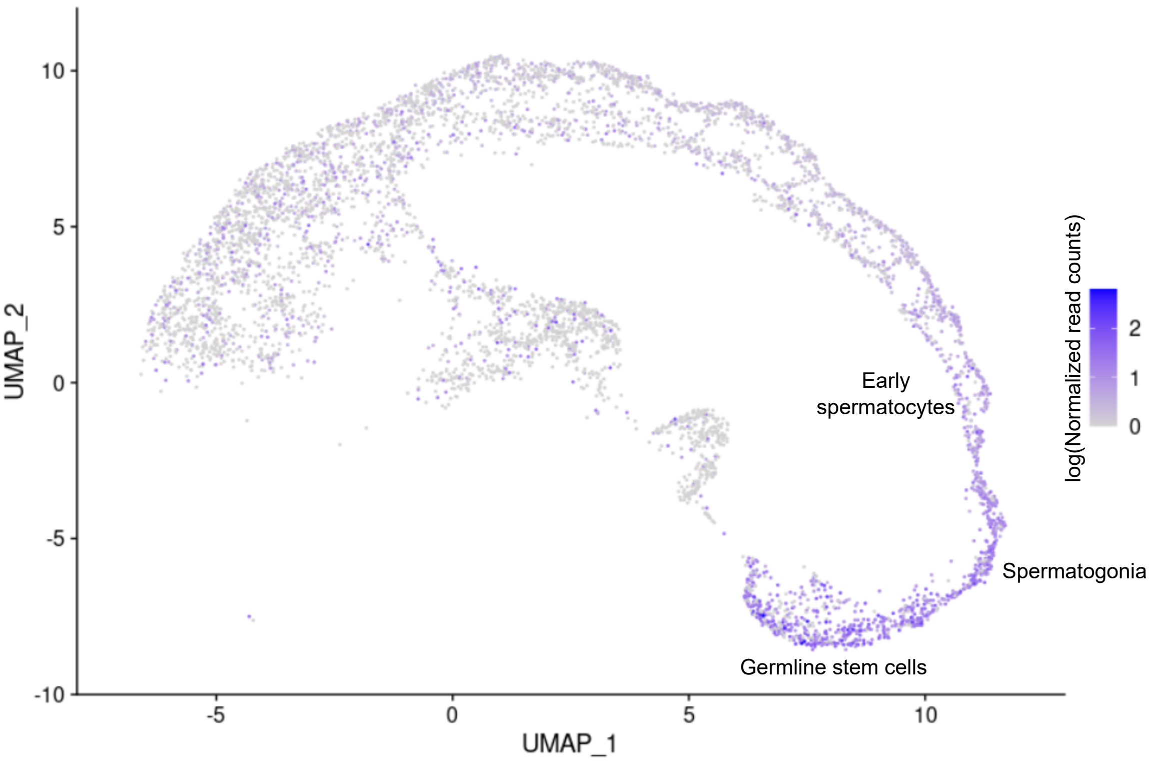


**Supplementary Figure 14:** Expression of c(3)G in testes single cell RNA-seq data for D. miranda. UMAP projection of testes cell types. Relevant cell types are labeled.

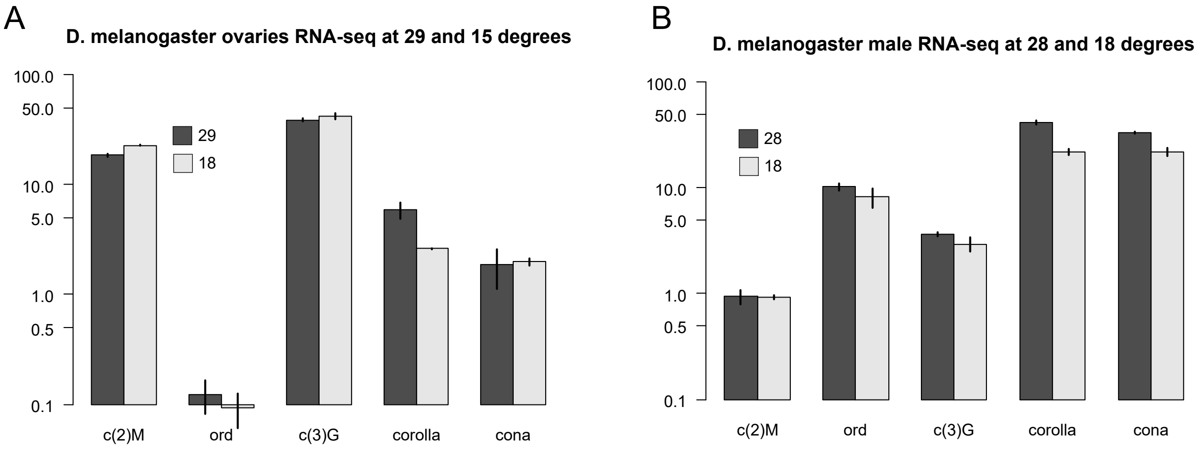


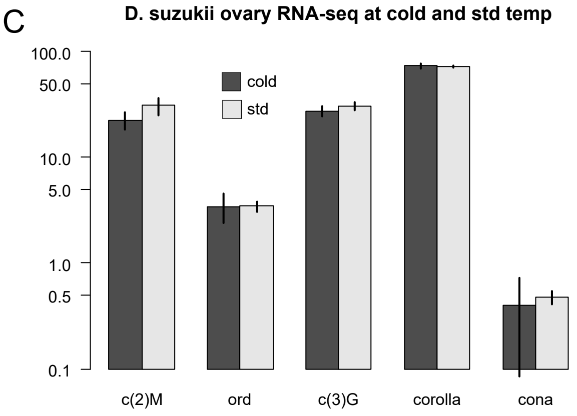


**Supplementary Figure 15:** Expression of SC genes in different temperatures in D. melanogaster ovaries (A), D melanogaster males whole bodies (B), and D. suzukii testes (C).

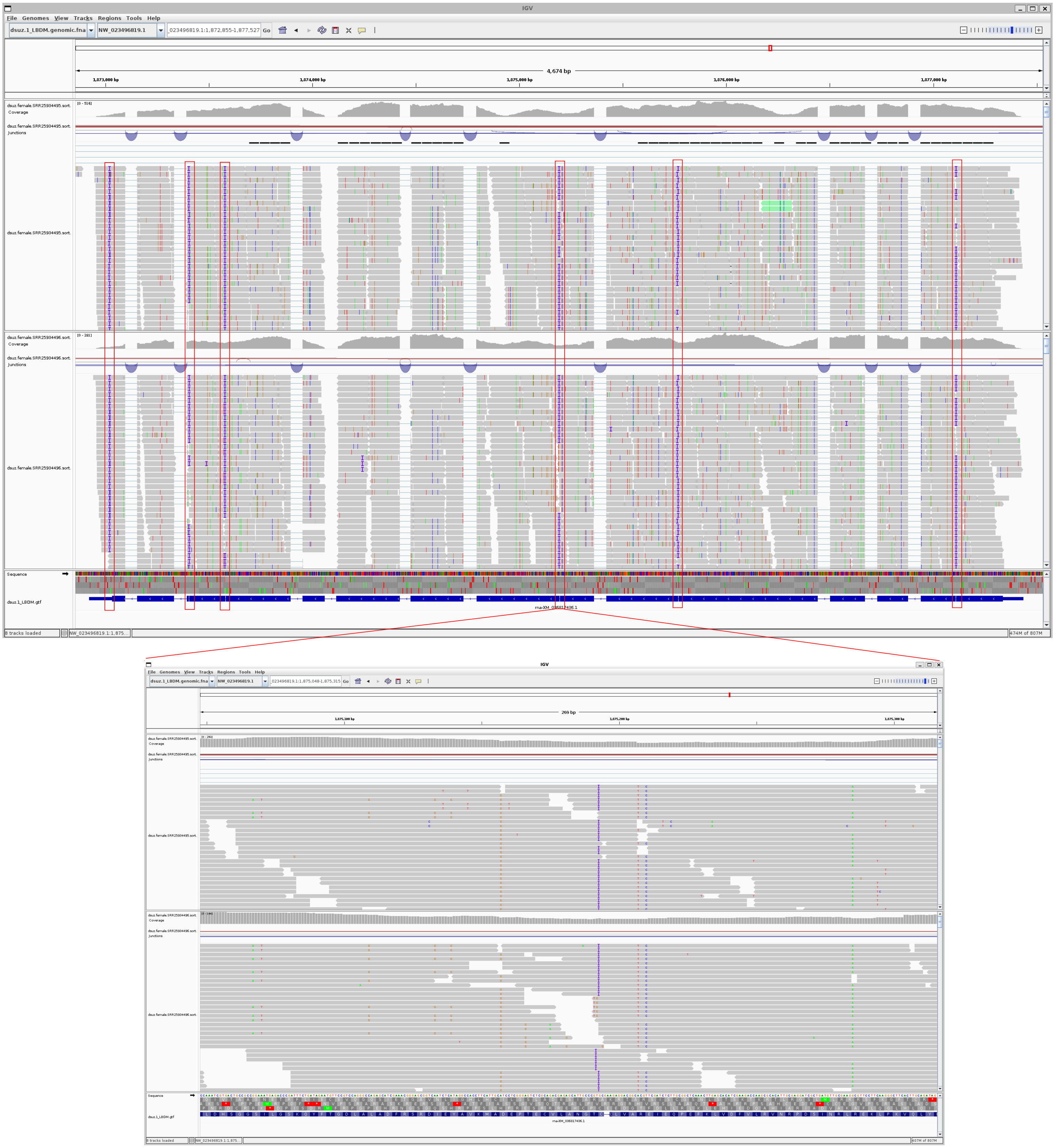


**Supplementary Figure 16:** Repeated indels at homopolymer tracks in the Refseq D. suzukii genome cause short introns in exons in NCBI annotations.
